# Identification of deleterious single nucleotide polymorphism (SNP)s on the human *TBX5* gene & prediction of their structural & functional consequences: An *in silico* approach

**DOI:** 10.1101/2020.05.16.099648

**Authors:** A M U B Mahfuz, Md. Arif Khan

## Abstract

T-box transcription factor 5 (*TBX5*) gene encodes the transcription factor TBX5, which plays a crucial role in developing the heart and upper limbs. Damaging single nucleotide variants in this gene alters the protein structure, disturbs the functions of TBX5, and ultimately causes Holt-Oram Syndrome (HOS). By analyzing available single nucleotide polymorphism information in the dbSNP database, this study was designed to identify the most deleterious *TBX5* SNPs through *silico* approaches and predict their structural and functional consequences.

Fifty-eight missense substitutions were found damaging by sequences homology-based tools like SIFT and PROVEAN and structure homology-based tools like PolyPhen-2. These SNPs were further scrutinized by various disease association meta-predictor tools. Additionally, the conservation profile of amino acid residues, their surface accessibility, stability, and structural integrity of the native protein upon mutations was assessed. From these analyses, finally 5 SNPs were detected as the most damaging ones [rs1565941579 (P85S), rs1269970792 (W121R), rs772248871 (V153D), rs769113870 (E208D), and rs1318021626 (I222N)]. Analyses of stop-loss, nonsense, UTR, and splice site SNPs were also conducted.

Through integrative bioinformatics analyses, this study has identified SNPs that are deleterious to the TBX5 protein structure and have the potential to cause HOS. Further wet-lab experiments can validate these findings.

## 1. Introduction

The heart is the first organ to develop in the human body, and numerous genes and their products orchestrate this process. Among these genes, *TBX5* is an important one. It is a member of the T-box gene family, which encode transcription factors with a highly conserved DNA binding domain (T-box) of 180-200 amino acids [1]. Members of the T-box gene family have been found from metazoans to humankind, and in humans, more than 20 members of this gene family have been reported till now [2, 3]. Moreover, several human diseases are associated with mutations in the T-box genes, namely, DiGeorge Syndrome (*TBX1*), Ulnar-Mammary Syndrome (*TBX3*), Small Patella Syndrome (*TBX4*), Holt-Oram Syndrome (*TBX5*), Spondylocostal Dysostosis (*TBX6*), Cousin Syndrome (*TBX15*), Isolated ACTH Deficiency (*TBX19*), Congenital Heart Defects (*TBX20*), X-Linked Cleft Palate with or without Ankyloglossia (*TBX22*) [4].

According to the recent information retrieved from NCBI and UniProt databases, the *TBX5* gene is located on the long arm of chromosome 12 at position 24.21 (12q24.21), contains ten exons, and has three known transcript variants (1, 3, and 4). Transcript variants 1 and 4 encode a polypeptide of 518 amino acid residues and transcript variant 3 encodes a 468 (51-518) amino acid polypeptide with a shortened N-terminus. In the TBX5 transcription factor, the T-box spans from amino acid position 58 to 238. The binding site of TBX5 has been identified to be present in the upstream regions of various cardiac-expressed genes, such as cardiac α actin, atrial natriuretic factor, cardiac myosin heavy chain α, cardiac myosin heavy chain β, myosin light chain 1A, myosin light chain 1V and *Nkx2-5*. The consensus binding site for TBX5 corresponds to the first eight bases from one-half of the Brachyury binding 22 palindromic consensus sequence [5]. It was shown *in vitro* that TBX5, when in total length, binds to the target site mainly as a dimer and, when truncated, tends to bind mainly as a monomer with reduced dimer formation [5]. It was further showed *in vitro* that they could form a heterodimer complex with Nkx2-5, another essential transcription factor for cardiac development in the course of DNA binding [6]. TBX5 acts mainly as a transcriptional activator of genes responsible for cardiomyocyte maturation and cardiac septa formation during the early stages of heart development. It is necessary for the effective establishment of the cardiac conduction system [7]. TBX5 also plays a vital role in the very initiation of upper (fore) limbs [8], but the necessity of its presence during the limb outgrowth phase remains controversial [8, 9].

Holt-Oram Syndrome (HOS) is an autosomal dominantly inherited disease due to mutation in the *TBX5* gene. It is considered 100% penetrant with varying expressivity in the heart and upper extremity [10]. However, incomplete penetrance [11], lack of penetrance [12], somatic mosaicism [12], and probable germinal mosaicism [13] have also been reported for this disease. The majority of the patients suffer from this syndrome due to *de novo* mutations [14]. Cardiac presentations include atrial septal defect, ventricular septal defect, atrioventricular septal defect, pulmonary atresia/ stenosis, double outlet right ventricle, aortic valve insufficiency/ stenosis, tricuspid valve atresia, mitral valve abnormality, patent ductus arteriosus, tetralogy of pentalogy of Fallot, common arterial truncus, dextrocardia, right aortic arch, and other non-specified congenital heart diseases [15]. Upper limb anomalies may include the hypoplastic or absent thumb, triphalangeal thumb, accessory/ bifid thumb, aplasia/ hypoplasia of hand and fingers, syndactyly, radial with or without ulnar aplasia or hypoplasia, radio-ulnar synostosis, hypoplasia of humerus, aplasia of humerus, and clavicle abnormalities [15].

Nonsense mutation [16], missense mutation [17], frameshift mutation [16, 17], deletion [18], duplication [19] and splice site mutation [20,21,22] of *TBX5* gene have been reported to be responsible for HOS. It has been found that missense mutations tend to cause more serious cardiac anomalies than mutations that cause a truncated TBX5 protein (nonsense, frameshift, splice site point variants, intragenic deletions, or duplications) [12]. HOS is caused by heterozygous mutation. To the best of our knowledge, only one homozygous missense mutation in the *TBX5* gene has been reported until present but only with cardiac involvement (atrioventricular septal defect) and no skeletal involvement [23].

*TBX5* missense mutations are also associated with familial [24] and sporadic dilated cardiomyopathy [25], atrial fibrillation, and lone atrial fibrillation [26]. *Furthermore, TBX5* non-synonymous (missense and nonsense) mutations have also been reported in association with tetralogy of Fallot without skeletal abnormality [27], and a *TBX5* 3′UTR variant makes its carriers more prone to congenital heart diseases [28]. Considering the above-stated findings, this study was conducted to find out the most deleterious *TBX5* SNPs and analyze their structural and functional outcomes *in silico*.

## 2. Materials and Methods

The overall methodology adopted in this study is graphically presented in Figure 1.

**Fig. 1:**
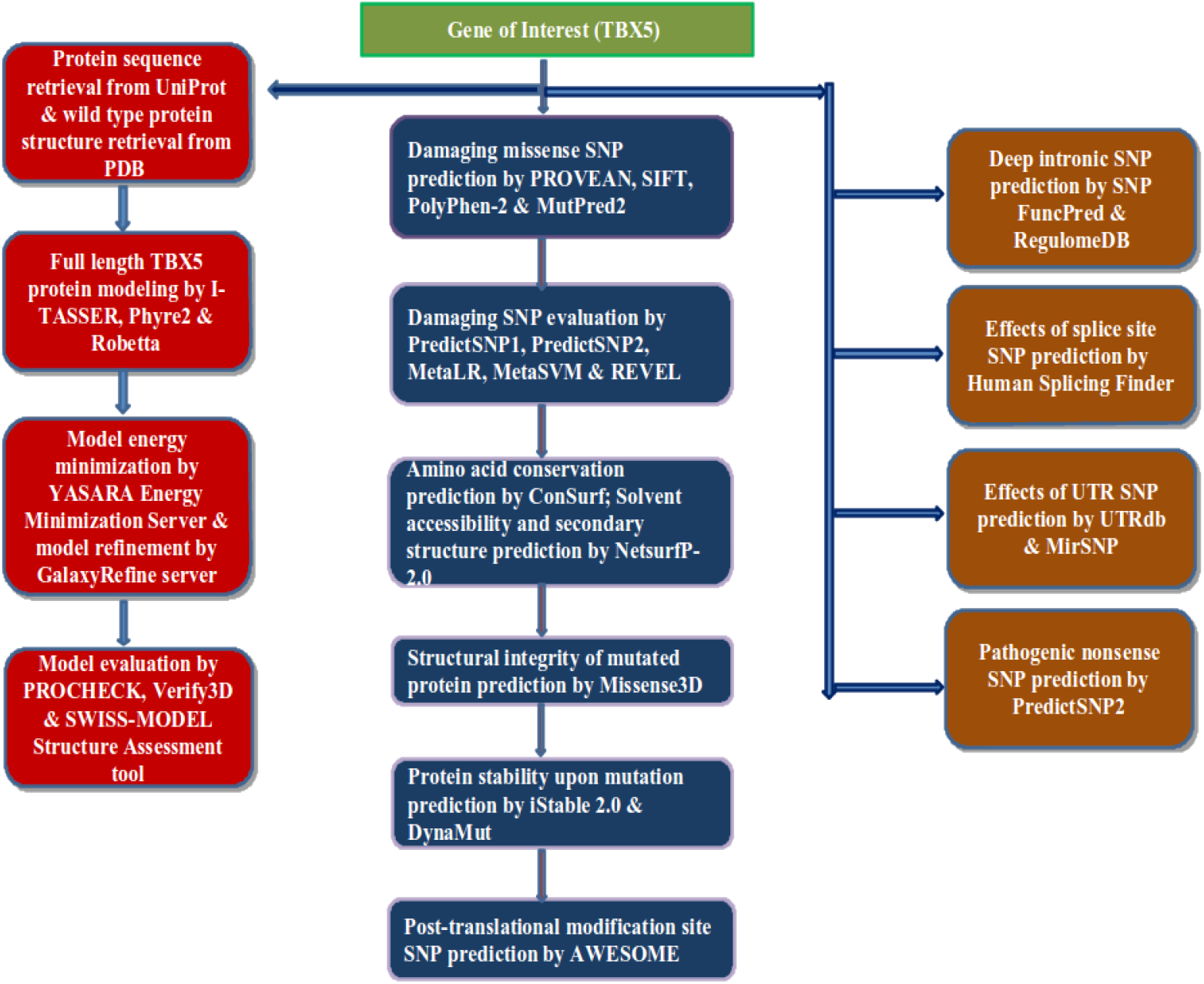
An overview of the steps followed to identify deleterious *TBX5* SNPs and analyze their structural and functional consequences.

### 2.1 Retrieval of Sequence and Wild Structure

Amino acid sequence of the human *TBX5* protein was downloaded in FASTA format from the UniProt database (UniProt ID: Q99593) (https://www.uniprot.org/uniprot/Q99593). Experimental structures of TBX5 were collected from Research Collaboratory for Structural Bioinformatics Protein Data Bank (RCSB PDB) (https://www.rcsb.org/).

### 2.2 Retrieval of SNPs

SNP information of the human *TBX5* gene was collected using NCBI Variation Viewer (https://www.ncbi.nlm.nih.gov/variation/view/). Nonsense, stop lost, splice acceptor, splice donor, and UTR SNPs of *TBX5* gene along with their rs (reference SNP) IDs were retrieved from Variation Viewer applying the following filters: dbSNP under Source database menu, single nucleotide variant under ‘Variant type’ menu, and nonsense (stop gained)/ stop lost/ splice acceptor variant/ splice donor variant/ 5 prime UTR variant/ 3 prime UTR variant under ‘Molecular consequence’ menu. In addition, information about missense SNPs was downloaded directly from the NCBI dbSNP database (https://www.ncbi.nlm.nih.gov/snp/).

### 2.3 Identification and Analysis of Damaging Missense SNPs

A total of 355 missense SNPs were retrieved from dbSNP.

#### 2.3.1 Prediction of Damaging Missense SNPs by Sequence-Homology Based Tools

The missense SNPs were first subjected to analysis by sequence-homology-based predictor PROVEAN [29]. PROVEAN predicts the effect of single or multiple amino acid substitutions or indel mutations on the functions of a protein. In PROVEAN, BLAST hits with more than 75% global sequence identity are clustered first. The top 30 clusters constitute the supporting sequence set, and a delta alignment score is computed for every supporting sequence. These scores are averaged within and across clusters to obtain the final PROVEAN score. An amino acid substitution is predicted ‘Deleterious’ if the PROVEAN score is below or equal to a predefined threshold value of -2.5 and is considered ‘Neutral’ if the PROVEAN score is above this threshold value [29]. In this study, a list of *TBX5* gene SNPs defined by their chromosomal positions, reference alleles, and variant alleles was used as input to the PROVEAN server.

PROVEAN also provides scores from SIFT analysis, another prediction tool. SIFT (Sorting Intolerant From Tolerant) presumes that essential amino acids are well conserved in a protein family, and a replacement in a conserved zone is predicted as ‘Damaging’ [30]. SIFT scores lie between 0 and 1, and amino acid substitutions with a score of ≤0.05 are predicted to alter protein function, and hence, are considered ‘Damaging’ [31].

85 SNPs of the *TBX5* gene were predicted as ‘Deleterious’ by PROVEAN, and 236 SNPs were predicted as ‘Damaging’ by SIFT. 80 SNPs were predicted as harmful by both tools.

#### 2.3.2 Prediction of Damaging Missense SNPs by Structure-Homology Based Tool PolyPhen-2

PolyPhen-2 (Polymorphism Phenotyping version 2) prediction is made on eight sequence-based and three structure-based predictive features. Two pairs of datasets, viz. HumDiv and HumVar were recruited to train and test two PolyPhen-2 models. PolyPhen-2 computes the Naive Bayes posterior probability of a given mutation is being damaging and estimates the false-positive rate and the true-positive rate. It classifies SNPs as ‘Probably Damaging,’ ‘Possibly Damaging,’ or ‘Benign’ based on the false-positive rate (FPR) thresholds [32]. The 80 SNPs that were predicted harmful by both PROVEAN and SIFT were next analyzed by both PolyPhen-2 HumDiv and HumVar using the batch query option. A list of SNPs defined by chromosome number, chromosomal position, reference allele, and variant allele was used as input. 58 SNPs were predicted as ‘Probably Damaging’ by either HumDiv or HumVar, and these SNPs were considered high fidelity in disease association.

#### 2.3.3 Prediction of Pathogenic Amino Acid Substitutions by MutPred2

MutPred2 integrates genetic and molecular information to predict pathogenicity caused by an amino acid substitution in the query protein [33]. The cutoff value for predicted pathogenicity is 0.5, and the higher the score, the greater the probability of that amino acid substitution’s disease association. TBX5 protein sequence in FASTA format, and wild type and substituted amino acids of 58 high-confidence deleterious SNPs with their respective positions were submitted to the MutPred2 server using the default P-value threshold (0.05).

#### 2.3.4 Scrutinization of Predicted Pathogenicity by Meta-predictors

Several meta-predictors have been developed for predicting disease-causing SNPs in recent years. Such meta-predictors employ various single pathogenicity prediction tools and provide a consensus score gained from different ensemble methods. It is well established that ensemble methods render improved predictive performance than any of its constituent algorithms alone. We decided to crosscheck the predicted 58 pathogenic SNPs against 5 such meta-predictors, namely, i) PredictSNP2, ii) PredictSNP1, iii) MetaLR, iv) MetaSVM and v) REVEL.

PredictSNP2 generates a consensus score from the prediction of five tools-CADD, DANN, FATHMM, FunSeq2, and GWAVA [34]. The input to PredictSNP2 was similar to that used for PolyPhen-2. After analysis by PredictSNP2, the SNPs were sent for analysis by PredicteSNP1. PredictSNP1 has incorporated six individual predictors-MAPP, PhD-SNP, PolyPhen-1, PolyPhen-2, SIFT, and SNAP [35]. MetaSVM and MetaLR are two meta-classifiers developed by the dbNSFP (database for non-synonymous SNPs’ functional predictions) project. They calculate ensemble scores based on the prediction of deleteriousness by 10 component tools and the maximum frequency observed in the 1000 genomes project populations [36]. The difference between them lies in their adopted approaches-one uses the Support Vector Machine (MetaSVM) algorithm, and the other uses the Logistic Regression (MetaLR) algorithm. They were both accessed through Ensembl Variant Effect Predictor (VEP) (http://www.ensembl.org/Tools/VEP). rsID (Reference SNP ID number)s of the SNPs were used as inputs and the reference genome chosen was GRCh37. REVEL (Rare Exome Variant Ensemble Learner) is a meta-predictor that integrates 13 tools to predict the probability of disease causation of a missense SNP. It is particularly efficient in differentiating pathogenic from rare neutral variants with allele frequencies <0.5% [37]. REVEL scores range between 0 and 1, and the threshold score for pathogenic variants is 0.5. REVEL scores in this study were obtained alongside MetaSVM and MetaLR scores using the Ensembl VEP.

#### 2.3.5 Prediction of Conserved Amino Acid Residues by ConSurf

structural and functional importance of an amino acid residue in a protein is often strongly related to its level of conservation. Consurf ranks evolutionary conservation status of amino acid or nucleic acid positions in a protein or DNA/ RNA molecule, respectively [38]. The continuous conservation scores of residues are split into an integer scale of 1 to 9 to depict a color scheme. The most variable positions (grade 1) are colored turquoise, intermediate conserved positions (grade 5) are colored white, and the most conserved positions (grade 9) are colored maroon. TBX5 protein sequence in FASTA format served as the input.

#### 2.3.6 Solvent Accessible Surface Area (SASA) Prediction by NetSurfP-2.0

Knowing the solvent accessibility of amino acid residues is essential to identify the interaction interfaces or active sites in a fully folded protein. Amino acid substitutions in such sites may bring change in binding affinity [39] or disturb catalytic activity if the protein is an enzyme [40]. NetSurfP-2.0 predicted surface accessibility of TBX5 residues. It accepts a protein sequence in FASTA format as input and recruits deep neural networks that have been trained on crystal protein structures [41].

#### 2.3.7 Prediction of Change in Stability of the Mutated Protein by Consensus Prediction

Amino acid substitutions caused by missense SNPs can significantly alter the folding free energy. Missense SNPs decreasing the stability of the native protein can affect the functions of the protein and ultimately result in diseases [42]. Missense SNPs causing destabilizing proteins are associated with myopathy, von Willebrand disease, retinitis pigmentosa, hemophagocytic lymphohistiocytosis, and prion diseases. However, stabilizing missense SNPs can also cause disease [42]. In our study, changes in TBX5 protein stability due to missense SNPs were predicted by iStable 2.0 [43]. iStable 2.0 results are derived from 11 protein stability prediction tools [43]. For amino acid substitutions up to F232V in TBX5, chain A of the TBX5 dimer crystal structure containing 1-239 residues (PDB ID: 5BQD) was used as the native structure. Due to unavailability of an experimental structure of the rest of the substitutions, they were analyzed using the protein’s FASTA sequence.

#### 2.3.8 Prediction of the Effects of Amino Acid Substitutions on the Structural Integrity of TBX5 Protein by Missense3D and Normal Mode Analysis

To analyze the impacts of amino acid substitutions on TBX5 three-dimensional structure, the Missense3D server was utilized. Missense3D considers 16 structural features to differentiate disease-associated SNPs from the neutral ones [45]. For substitutions up to F232V, the ‘Position on Protein Sequence’ input option was chosen, and the UniProt ID (Q99593) and PDB code & chain ID (5BQD; A) along with the substitutions were used as input. For the rest of the substitutions, the energy minimized model generated by Robetta was used [see 2.8]. The mutated protein structures due to SNPs were later subjected to normal mode analysis by DynaMut [44]. DynaMut calculates probable consequences of an amino acid replacement on the stability of a submitted protein from vibrational entropy changes. Normal mode analysis is a widely applied tool for protein structure study [44]. Input was similar to Missense3D.

#### 2.3.9 Molecular Dynamics Simulation

Molecular dynamics (MD) simulations of five mutant structures distilled from the consensus of Missense3D, DynaMut and iStable 2.0 [see 3.2.7], and native protein structure (PDB ID:5BQD, chain A) were conducted by SimpleRun module of PlayMolecule suite [46]. The proteins were initially prepared for MD by the ProteinPrepare module (https://www.playmolecule.org/proteinPrepare/). System for MD simulations were set by the SystemBuilder module (https://www.playmolecule.org/SystemBuilder/). To mimic physiological environment, a solvation box (90×90×90 Å) with pH 7.4 and 0.9% (0.154 M) NaCl concentration was chosen. Selected solvation model was TIP3P. MD simulation was continued for 25 ns for each protein structure by applying AMBER (Assisted Model Building with Energy Refinement) force field. The trajectories were saved at 100 pico second intervals to calculate RMSD and RMSF.

#### 2.3.10 Prediction of Post-translational Modification Site Missense SNPs by AWESOME

Protein post-translational modification (PTM)s occur at the time of or after protein synthesis, and such modifications of protein structures are mediated by different enzymes or covalent bonds with other structures. SNPs in the PTM sites can result in diseases [47]. SNPs in the putative PTM sites of the TBX5 protein were detected by AWESOME [47]. AWESOME predicts the PTM site SNPs based on the evaluation of 20 PTM site predictors. Gene’s name was used as input to AWESOME.

### 2.4 Analysis of Nonsense and Stop Lost SNPs by PredictSNP2

31 nonsense (stop gained) and one-stop lost SNPs were retrieved from NCBI Variation Viewer. Among the 31 nonsense SNPs, 29 are already annotated as ‘Pathogenic’ in the ClinVar database (https://www.ncbi.nlm.nih.gov/clinvar/). Rest 2 nonsense SNPs, rs1393693495 (W514Ter) and rs1555226308 (Q156Ter), and 1 stop lost SNP, rs1394777873 (Ter519Q) were analyzed by PredictSNP2 [35].

### 2.5 Prediction of the Effects of UTR SNPs by UTRdb & MirSNP

Untranslated region (UTR)s play a pivotal role in the regulation of gene expression post-transcriptionally. They ascertain nucleo-cytoplasmic transport of mRNA, maintain translational efficiency, subcellular localization, and RNA stability [48]. The essential regulators in the 5′-UTR are Kozak consensus sequence (ACCAUGG) [Shine-Dalgarno consensus sequence (AGGAGG) in prokaryotes], CpG (5’—C— phosphate—G—3’) sites, uORFs (upstream open reading frames), IRESs (Internal Ribosome Entry Sites), and RBP (Ribosome-binding protein) binding sites. 3′-UTR length, RBP binding sites, and miRNA binding sites are the critical players in the 3′-UTR [49]. Single nucleotide variants in these regions have been found to be associated with many diseases like thalassemia intermedia, atrial septal defect [50], campomelic dysplasia [51], hereditary chronic pancreatitis [52], Marie Unna hereditary hypotrichosis [53], Charcot-Marie-Tooth disease [54], cerebral amyloid angiopathy [55], to name a few. So, it is essential to identify potentially harmful SNPs in the 3′ & 5′-UTR. UTR SNPs that were predicted to affect transcriptional motifs have been retrieved from the UTRsite database [56]. ‘UTRef’ (NCBI RefSeq transcripts) as the searching database and ‘Gene Name’ as the accession type were chosen during retrieval.

MicroRNA (miRNA)s also have critical roles in post-transcriptional regulation of gene expression through mRNA cleavage or post-translational repression [57]. miRNAs are small non-coding RNAs having about 22 nucleotides. miRNA seed region (positions 2-8 on miRNA) is the most crucial sequence in finding the complementary sequence on mRNA and subsequently binding with it. SNPs in the miRNA target sites in the 3′ UTR may create or delete a miRNA target or change the binding efficiency between miRNA and mRNA. The creation or deletion of a miRNA target site may also affect binding in its neighboring miRNA target sequences. Such SNPs have been reported in several diseases, including cancers, psychiatric illnesses, cardiomyopathy, asthma, and Parkinson’s disease [58, 59]. In our study, SNPs in the predicted miRNA target sites were identified using the MirSNP server [59]. Gene name was used as input to MirSNP.

### 2.6 Prediction of the Effects of Splice Site SNPs by HSF (Human Splicing Finder)

RNA splicing is necessary for converting precursor mRNA to a mature mRNA ready for translation. Various *cis*-regulatory elements and *trans-*acting elements maintain this splicing process. *Cis*-regulatory elements include 5′ and 3′ consensus splice site sequences in the exon-intron boundaries and branch point and polypyrimidine tract sequences. *Trans-*acting elements include spliceosomal small nuclear RNAs and proteins and various other splicing repressors and activators, many of which are still unknown [60, 61]. Mutations that hamper the controlled process of splicing lead to a good number of diseases. Around 9% of all the pathogenic mutations recorded in The Human Gene Mutation Database are splicing ones [60, 61]. SNPs in the splice donor and acceptor sites may break an established splice site or create a new splice site. They may also activate cryptic sequences (sequences similar to splice sites) or affect splicing enhancers and silencers. They may even intercept the binding of spliceosomal components by making a conformational change in the mRNA secondary structure [60]. In this study, seven splice donor single nucleotide variants and seven splice acceptor single nucleotide variants were retrieved from dbSNP using NCBI Variation Viewer. Among the seven donor variants, two are annotated as ‘Likely-pathogenic’. Among the seven acceptor variants, one is annotated as ‘Pathogenic’ and three are annotated as ‘Likely-pathogenic’ in the ClinVar database. Excluding them, the rest 8 SNPs were analyzed employing Human Splicing Finder (HSF) [62]. For our analysis, ‘Analyze a sequence’ and ‘Paste your own sequence’ options were chosen, and individual variants with a flanking sequence of 100 nucleotides in both 5′ and 3′ directions were used as inputs.

### 2.7 Prediction of the Effects of Deep Functional Intronic SNPs

Intronic regions located >100 base pairs away from the exon-intron borders are known as deep intronic regions [63]. Although they have been overlooked for a long time, they are now considered necessary, and mutations in these regions have been found in association with more than 75 diseases. Furthermore, deep intronic point mutations, deletions, and insertions are held accountable for these disease causations, with the first one being the most frequent [63].

The rs (reference SNP) IDs of all the *TBX5* intron variants were submitted to SNP Function Prediction (FuncPred) using default settings [64]. 19 SNPs were predicted by FuncPred to affect TF binding sites and were next analyzed by RegulomeDB 2.0 server. RegulomeDB 2.0 is an annotated database of SNPs in the human intergenic regions that have either established or predicted regulatory roles [65].

### 2.8 3D Structure Prediction of Full TBX5 Protein

Currently, there are four experimental structures of TBX5 (2X6U, 2X6V, 4SOH & 5BQD) deposited in the Protein Data Bank (PDB). Nevertheless, all of them cover less than half of the total protein. So, we decided to predict the structure of the whole TBX5 protein by three reliable protein structure prediction tools, namely, I-TASSER [66], Phyre2 [67], and Robetta [68]. Energy minimization of the generated TBX5 models was achieved by applying the YASARA force field. YASARA force field is a combination of AMBER all-atom force field and multi-dimensional knowledge-based torsional potentials [69]. The energy minimized models were further refined by the GalaxyRefine server (http://galaxy.seoklab.org/cgi-bin/submit.cgi?type=REFINE).

### 2.9 Evaluation of TBX5 3D Models

Quality of the three predicted TBX5 models was evaluated by PROCHECK [70], Verify 3D [71], and SWISS-MODEL Structure Assessment Tool [72]. PROCHECK examines specific stereochemical and geometrical properties of the query protein structure and compares them against a set of ideal values of these properties obtained from well-refined high-resolution protein structures. It assesses residue-by-residue geometry and the overall structural geometry of the query structure [70]. A Ramachandran plot of *φ*-*ψ* torsion angles generated by PROCHECK is the most helpful quality indicator, and a good quality model is expected to have over 90% residues in the most favored regions. Verify 3D checks the agreement between a protein tertiary structure and its primary structure [71]. SWISS-MODEL Structure Assessment Tool provides assessments from the MolProbity and QMEAN algorithms. These algorithms evaluate a model quality at both global (i.e., for the entire structure) and local (i.e., per residue) level [72, 73]. QMEAN Z-score is an indicator of the “degree of nativeness” of a model [74].

## 3. Results

### 3.1 SNP Information Retrieval

Searching the NCBI Variation Viewer using dbSNP as the ‘source database’ and single nucleotide variant (SNV) as the ‘variant type’, returned 14413 rsIDs associated with the *TBX5* gene as single nucleotide variants. Three hundred fifty-five rsIDs were found to be missense variants, 30 rsIDs to be nonsense (stop gained) variants, one rsID to be ‘stop lost’ variant, 210 rsIDs to be synonymous variants, 11300 rsIDs to be intron variants, 821 rsIDs to be non-coding transcript variants, seven rsIDs to be splice acceptor and seven rsIDs to be splice donor variants, 689 rsIDs to be five prime UTR variants, and 345 rsIDs to be three prime UTR variants. Figure 2 is a pie chart showing different types of SNP. On further inspection, 404 missense variants with 355 rsIDs (3 nonsense variants are included in rs377649723, rs765204502, and rs903933027, and 11 synonymous variants are included in rs186780790, rs377532269, rs533581420, rs374600913, rs753688559, rs762204624, rs771883815, rs777853147, rs972633602, rs973631007, and rs769107196) and 28 nonsense variants with 28 rs IDs (27 stop gained and 1 stop lost-rs1394777873) were found. The reason behind a single rsID having more than one variant is that these variants were reported on the same chromosomal location. As per the NCBI rule, only one rsID is assigned for a specific chromosomal location [75].

**Fig. 2:**
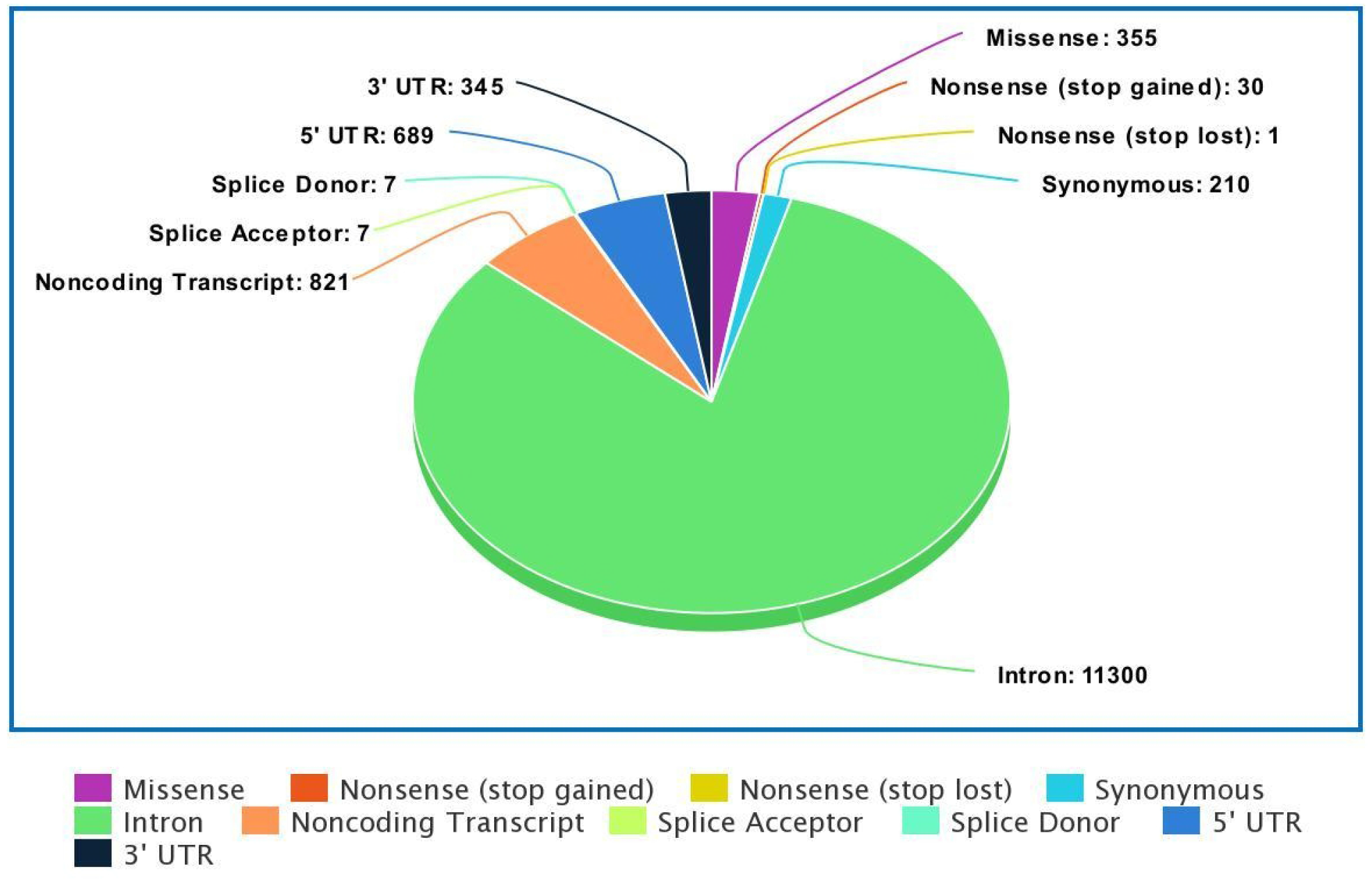
Pie-chart displaying the distribution of SNPs on the *TBX5* gene.

### 3.2 Identification and Analysis of Damaging Missense SNPs

#### 3.2.1 Prediction of Missense SNPs by Sequence-Homology Based Tools PROVEAN & SIFT

The retrieved 404 missense SNPs were analyzed first using PROVEAN. Out of 404 submitted substitutions, PROVEAN predicted 85 as ‘Deleterious’ and 301 as ‘Neutral’, and SIFT predicted 236 as ‘Damaging’ and 150 as ‘Tolerated’. 242 substitutions were predicted as ‘Deleterious’ or ‘Damaging’ by either PROVEAN or SIFT, respectively, and 80 substitutions were predicted as ‘Deleterious’ or ‘Damaging’ by both tools. Results for rs571328934, rs755664220, rs756902625, rs763178387, rs769107196, rs769895888, rs777370158, rs907398622, rs945043550, rs1263504354, rs1407437475, and rs1489969947 could not be predicted by PROVEAN.

#### 3.2.2 Prediction of Missense SNPs by Structure-Homology Based Tool PolyPhen-2

For high fidelity, the 80 SNPs predicted by PROVEAN and SIFT as ‘Deleterious’/ ‘Damaging’ were next analyzed by PolyPhen-2. Both HumDiv and HumVar dataset trained PolyPhen-2 were used. The HumDiv model predicted 58 substitutions as ‘probably damaging’ (Supplementary Table 1), 16 substitutions as ‘possibly damaging’, and 6 substitutions as ‘benign’. The HumVar model predicted 56 substitutions as ‘probably damaging’ (Supplementary Table 1), ten substitutions as ‘possibly damaging’, and 14 substitutions as ‘benign’. The 58 SNPs that were predicted as ‘probably damaging’ by HumDiv model were also predicted as ‘probably damaging by HumVar model except for rs190881877 (V205F) and rs944423586 (P337H) which were predicted as ‘possibly damaging’. rs1411518530 is associated with two nucleotide variants (G>C, T), but they encode the same amino acid (D110E) and are considered a single variant in the subsequent evaluations. Predictions by PROVEAN, SIFT, and PolyPhen-2 are available in Supplementary Table 1.

#### 3.2.3 Prediction of Pathogenic Amino Acid Substitutions by MutPred2

The threshold score of pathogenicity prediction by MutPred2 is 0.5, and a substitution having a MutPred2 score ≥0.8 can be considered highly likely to be pathogenic. All but two (K226R and P337H) of the 58 high-confidence deleterious SNPs have a MutPred2 prediction score ≥of 0.5. Supplementary Table 2 provides MutPred2 outcomes.

#### 3.2.4 Scrutinization of Predicted Pathogenicity by Meta-predictors

The pathogenicity of the aforementioned substitutions was further examined by five deleteriousness meta-predictors whose predictions are based on various ensemble methods. Forty-eight substitutions were predicted ‘Damaging’ by all the meta-servers, and the rest were predicted ‘Damaging’ by at least two meta-servers. The results are shown in Table 1.

**Table 1:** Prediction of amino acid substitution and disease relation by five meta servers.

| AAS | PredictSNP 1 |  | PredictSNP 2 |  | MetaLR |  | MetaSVM |  | REVEL |  |
| --- | --- | --- | --- | --- | --- | --- | --- | --- | --- | --- |
|  | Predict<br>ion | Expected<br>Accurac<br>y | Predictio<br>n | Expected<br>Accurac<br>y | Predictio<br>n | Sco<br>re | Predictio<br>n | Sc<br>ore | Predictio<br>n | Scor<br>e |
| L58P | D<sup>†</sup> | 0.87 | D | 0.87 | D | 0.93 | D | 1.08 | D | 0.98 |
| T72K | D | 0.87 | D | 0.87 | D | 0.88 | D | 1.02 | D | 0.94 |
| T72M | D | 0.87 | D | 0.87 | D | 0.83 | D | 0.86 | D | 0.95 |
| G80R | D | 0.87 | D | 0.87 | D | 0.94 | D | 1.10 | D | 0.98 |
| F84L | D | 0.72 | D | 0.87 | D | 0.91 | D | 1.06 | D | 0.97 |
| P85L | D | 0.87 | D | 0.87 | D | 0.96 | D | 1.09 | D | 0.96 |
| P85S | D | 0.87 | D | 0.87 | D | 0.96 | D | 1.10 | D | 0.97 |
| V91A | D | 0.65 | D | 0.87 | D | 0.78 | D | 0.62 | D | 0.73 |
| G93S | D | 0.87 | D | 0.87 | D | 0.96 | D | 1.10 | D | 0.97 |
| G93D | D | 0.87 | D | 0.87 | D | 0.96 | D | 1.10 | D | 0.94 |
| G93V | D | 0.61 | D | 0.87 | D | 0.96 | D | 1.10 | D | 0.94 |
| P96S | N<sup>§</sup> | 0.63 | D | 0.87 | D | 0.86 | D | 0.87 | D | 0.83 |
| Y100C | D | 0.87 | D | 0.87 | D | 0.96 | D | 1.10 | D | 0.98 |
| Y100F | D | 0.76 | D | 0.82 | D | 0.93 | D | 1.08 | D | 0.93 |
| M104T | D | 0.65 | D | 0.87 | D | 0.86 | D | 0.93 | D | 0.93 |
| D105N | D | 0.87 | D | 0.87 | D | 0.89 | D | 1.01 | D | 0.87 |
| V107E | D | 0.87 | D | 0.82 | D | 0.76 | D | 0.67 | D | 0.92 |
| P108T | D | 0.72 | D | 0.87 | D | 0.86 | D | 0.92 | D | 0.88 |
| D110E | D | 0.87 | N | 0.65 | D | 0.83 | D | 0.76 | D | 0.72 |
| D110N | D | 0.87 | D | 0.87 | D | 0.86 | D | 0.91 | D | 0.77 |
| D111Y | D | 0.76 | D | 0.87 | D | 0.82 | D | 0.79 | D | 0.87 |
| R113I | D | 0.87 | D | 0.82 | D | 0.89 | D | 1.00 | D | 0.95 |
| R113K | D | 0.61 | D | 0.87 | D | 0.83 | D | 0.84 | D | 0.92 |
| Y114H | D | 0.87 | D | 0.87 | D | 0.85 | D | 0.89 | D | 0.97 |
| W121R | D | 0.87 | D | 0.87 | D | 0.97 | D | 1.08 | D | 0.98 |
| G125R | D | 0.87 | D | 0.87 | D | 0.89 | D | 1.00 | D | 0.90 |
| G133C | N | 0.63 | D | 0.87 | D | 0.76 | D | 0.60 | D | 0.68 |
| L135R | D | 0.87 | D | 0.87 | D | 0.78 | D | 0.72 | D | 0.87 |
| V137A | D | 0.51 | D | 0.87 | D | 0.79 | D | 0.74 | D | 0.89 |
| D140E | N | 0.65 | D | 0.87 | D | 0.79 | D | 0.65 | D | 0.74 |
| P142S | D | 0.87 | D | 0.87 | D | 0.84 | D | 0.90 | D | 0.76 |
| A143T | D | 0.55 | D | 0.87 | D | 0.81 | D | 0.73 | D | 0.59 |
| G145W | D | 0.87 | D | 0.87 | D | 0.98 | D | 1.04 | D | 0.89 |
| V153D | D | 0.87 | D | 0.82 | D | 0.89 | D | 1.00 | D | 0.95 |
| L160F | D | 0.87 | D | 0.87 | D | 0.89 | D | 1.02 | D | 0.88 |
| H170L | D | 0.72 | N | 0.63 | D | 0.78 | D | 0.75 | D | 0.96 |
| N174D | D | 0.87 | D | 0.87 | D | 0.87 | D | 0.99 | D | 0.81 |
| V186G | D | 0.87 | D | 0.87 | D | 0.87 | D | 0.93 | D | 0.94 |
| V186M | D | 0.61 | D | 0.87 | D | 0.86 | D | 0.86 | D | 0.85 |
| D189Y | D | 0.87 | D | 0.87 | D | 0.84 | D | 0.80 | D | 0.68 |
| H204D | D | 0.87 | D | 0.87 | D | 0.80 | D | 0.69 | D | 0.88 |
| V205F | D | 0.72 | D | 0.87 | D | 0.80 | D | 0.70 | D | 0.75 |
| E208D | D | 0.51 | D | 0.87 | D | 0.84 | D | 0.80 | D | 0.71 |
| A213T | D | 0.55 | D | 0.87 | D | 0.85 | D | 0.71 | D | 0.72 |
| I222N | D | 0.87 | D | 0.87 | D | 0.89 | D | 0.94 | D | 0.90 |
| T223M | D | 0.87 | D | 0.87 | D | 0.91 | D | 1.05 | D | 0.96 |
| K226R | D | 0.76 | D | 0.87 | D | 0.92 | D | 1.01 | D | 0.88 |
| K226E | D | 0.87 | D | 0.87 | D | 0.93 | D | 1.10 | D | 0.96 |
| F232V | D | 0.87 | D | 0.82 | D | 0.92 | D | 1.08 | D | 0.95 |
| R237P | D | 0.87 | D | 0.87 | D | 0.91 | D | 1.06 | D | 0.95 |
| R237Q | D | 0.87 | D | 0.87 | D | 0.89 | D | 1.00 | D | 0.92 |
| R237W | D | 0.87 | D | 0.87 | D | 0.90 | D | 1.03 | D | 0.91 |
| S261C | D | 0.55 | D | 0.82 | D | 0.71 | D | 0.54 | D | 0.88 |
| C323Y | D | 0.72 | D | 0.87 | D | 0.85 | D | 0.64 | T | 0.47 |
| P337H | D | 0.61 | D | 0.87 | T<sup>‡</sup> | 0.20 | T | -0.66 | T | 0.20 |
| R375W | D | 0.76 | D | 0.87 | T | 0.33 | T | -0.32 | T | 0.38 |
| W401C | D | 0.72 | D | 0.82 | T | 0.33 | T | -0.36 | T | 0.44 |

AAS=Amino Acid Substitution, <sup>†</sup>D=Deleterious/Damaging/Disease, <sup>‡</sup>T=Tolerated, <sup>§</sup>N=Neutral

#### 3.2.5 Prediction of Conserved Amino Acid Residues by ConSurf and Solvent Accessibility of Residues and TBX5 Secondary Structure by NetSurfP-2.0

ConSurf provided position-specific amino acid conservation scores of TBX5 residues. It also ranked the amino acid residues in a color-coded scale of integers ranging from 1 to 9, where one indicates a highly variable and 9 indicates the most conserved residue. It was found that most of the substitutions are located in highly conserved positions. NetSurfP-2.0 predicted solvent accessibility (exposed or buried) of the amino acids and provided each residue’s relative and total accessible surface area. 37 substitutes were found to occur in buried residues, and 21 substitutes were found to be in exposed residues. Table 2 summarizes ConSurf and NetSurfP-2.0 outcomes. Figures 3 and 4 are graphical presentations of these results.

**Fig. 3:**
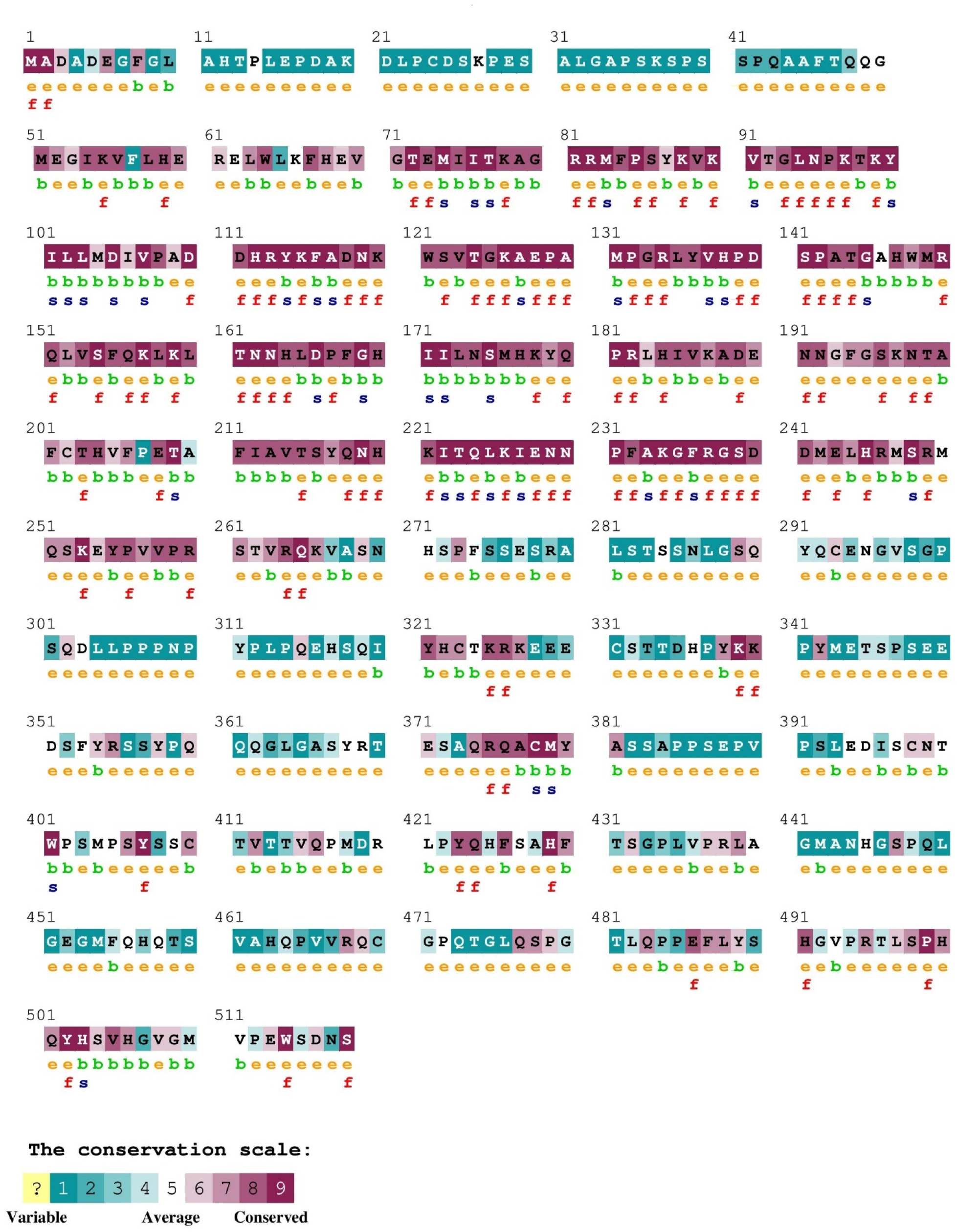
Detection of conserved amino acids by ConSurf. Here, e= an exposed residue according to the neural-network algorithm, b= a buried residue according to the neural-network algorithm, f= a predicted functional residue (highly conserved and exposed), s= a predicted structural residue (highly conserved and buried).

**Fig. 4:**
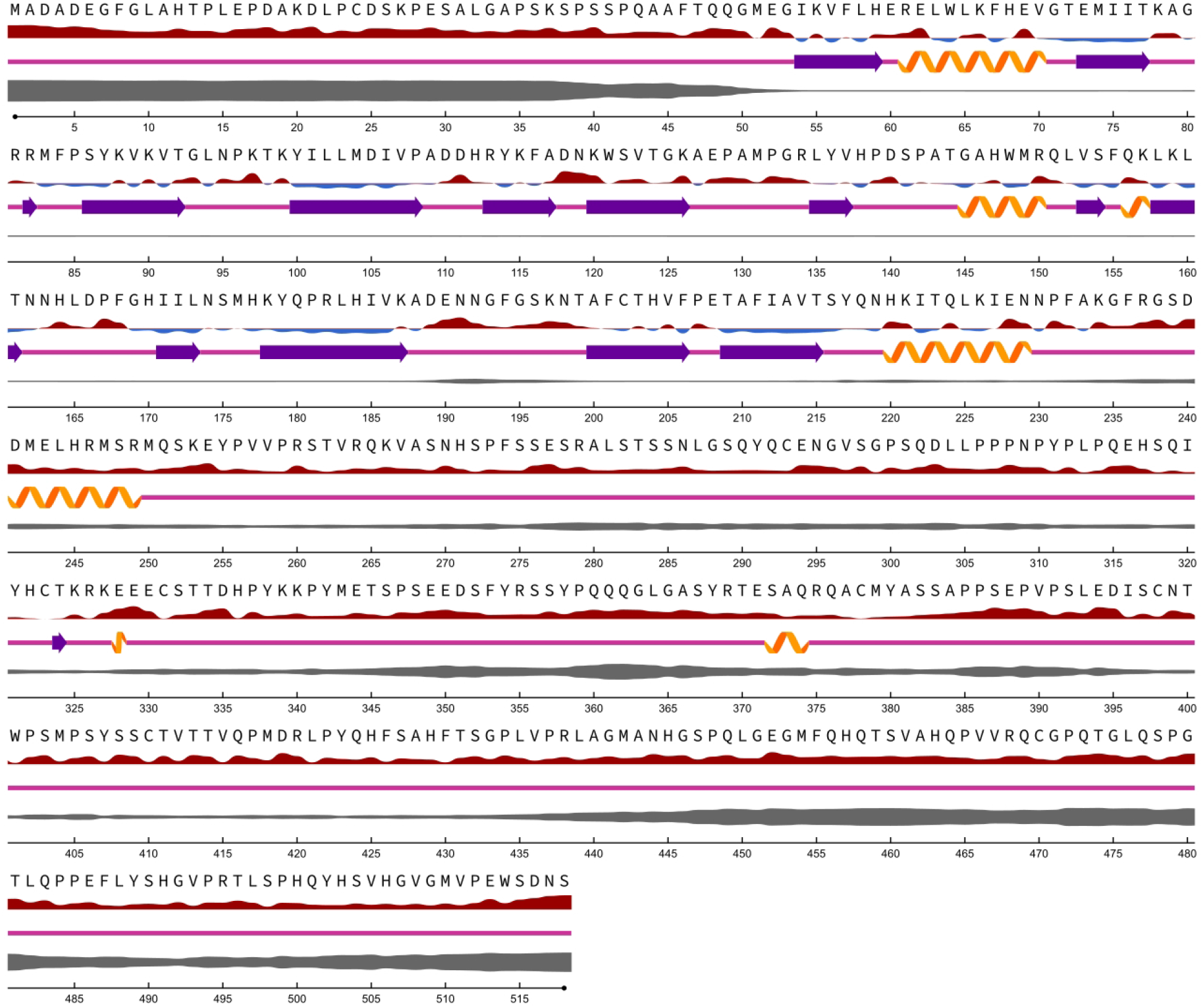
Solvent accessibility prediction of TBX5 residues by NetSurfP-2.0. Here, **Relative Surface Accessibility:** Red upward elevations are exposed residues and sky-blue low elevations are buried residues, thresholded at 25%, **Secondary Structure:** orange spiral=helix, indigo arrow=strand, and pink straight line=coil, **disorder:** swollen black line; thickness of line corresponds to the probability of disorder.

**Table 2:** Prediction by ConSurf and NetSurfP Webtools.

| ConSurf |  |  |  |  | NetSurfP-2.0 |  |  |
| --- | --- | --- | --- | --- | --- | --- | --- |
| Position | AA | Normalized Conservation Score | COLOR | RESIDUE VARIETY | Assigned Class | RSA<sup>†</sup> | ASA<sup>‡</sup> (Å) |
| 58 | L | -0.784 | 8 | T,L,M,H | Buried | 0.068 | 12.397 |
| 72 | T | -1.098 | 9 | T, S | Buried | 0.094 | 13.02 |
| 80 | G | -0.858 | 8 | N,H,G | Buried | 0.184 | 14.447 |
| 84 | F | -0.965 | 9 | C, F | Buried | 0.184 | 37.024 |
| 85 | P | -0.917 | 9 | G, V, P | Buried | 0.094 | 13.32 |
| 91 | V | -0.972 | 9 | L,X,V,C,M | Buried | 0.062 | 9.596 |
| 93 | G | -0.889 | 8 | K,G,R,X | Exposed | 0.526 | 41.429 |
| 96 | P | -0.715 | 8 | S,A,K,P,E | Exposed | 0.369 | 52.33 |
| 100 | Y | -1.095 | 9 | Y | Buried | 0.068 | 14.536 |
| 104 | M | -0.334 | 6 | M,V,I,A,T | Buried | 0.009 | 1.773 |
| 105 | D | -1.138 | 9 | D | Buried | 0.174 | 25.002 |
| 107 | V | -1.025 | 9 | I, V | Buried | 0.163 | 25.004 |
| 108 | P | -0.651 | 8 | A,T,S,Q,P | Buried | 0.218 | 31.002 |
| 110 | D | -1.138 | 9 | D | Exposed | 0.347 | 50.019 |
| 111 | D | -0.774 | 8 | E, D | Exposed | 0.653 | 94.129 |
| 113 | R | -1.137 | 9 | R | Exposed | 0.277 | 63.479 |
| 114 | Y | -1.095 | 9 | Y | Buried | 0.064 | 13.575 |
| 121 | W | -0.647 | 8 | W,R,C | Buried | 0.249 | 59.799 |
| 125 | G | -0.861 | 8 | G,P,S | Exposed | 0.293 | 23.038 |
| 133 | G | -0.794 | 8 | P,R,G,K | Exposed | 0.507 | 39.935 |
| 135 | L | -0.875 | 9 | M,L,P | Buried | 0.216 | 39.513 |
| 137 | V | -0.981 | 9 | P,I,V,L | Exposed | 0.292 | 44.873 |
| 140 | D | -0.944 | 9 | A,S,D | Exposed | 0.559 | 80.603 |
| 142 | P | -1.116 | 9 | P | Exposed | 0.297 | 42.08 |
| 143 | A | -0.875 | 8 | P,V,G,S,A | Buried | 0.223 | 24.613 |
| 145 | G | -0.901 | 9 | D,G,T | Buried | 0.042 | 3.271 |
| 153 | V | -0.761 | 8 | I, V, L | Buried | 0.041 | 6.232 |
| 160 | L | -0.81 | 8 | R,P,T,L | Buried | 0.034 | 6.174 |
| 170 | H | -0.889 | 8 | H,F,S,Q | Buried | 0.165 | 29.977 |
| 174 | N | -0.884 | 8 | A,G,S,N,R | Exposed | 0.279 | 40.854 |
| 186 | V | -0.825 | 8 | A,L,S,M,V,P | Buried | 0.046 | 7.095 |
| 189 | D | -0.802 | 8 | H,P,A,S,D,T | Exposed | 0.44 | 63.434 |
| 204 | H | -0.657 | 8 | H,R,F,Q,Y,A | Buried | 0.141 | 25.641 |
| 205 | V | -0.275 | 6 | W,A,S,L,V,M,I | Exposed | 0.44 | 67.687 |
| 208 | E | -0.846 | 8 | L,W,P,V,E | Exposed | 0.283 | 49.385 |
| 213 | A | -0.689 | 8 | G,T,S,A,P,M | Buried | 0.028 | 3.079 |
| 222 | I | -0.944 | 9 | I,V,N,W | Buried | 0.044 | 8.056 |
| 223 | T | -1.012 | 9 | T, Q, N | Buried | 0.159 | 22.092 |
| 226 | K | -1.021 | 9 | R, K | Buried | 0.128 | 26.428 |
| 232 | F | -0.808 | 8 | N,Y,F,L | Exposed | 0.4 | 80.298 |
| 237 | R | -1.099 | 9 | R | Exposed | 0.545 | 124.801 |
| 261 | S | -0.451 | 7 | N,D,S,T | Exposed | 0.424 | 49.711 |
| 323 | C | -0.522 | 7 | C,R,Y,L,S,G | Buried | 0.247 | 34.65 |
| 337 | P | 1.6 | 1 | G,T,D,L,S,Q,E,P,A,<br>H | Exposed | 0.578 | 81.958 |
| 375 | R | -0.72 | 8 | G,S,Q,R | Exposed | 0.412 | 94.327 |
| 401 | W | -1.02 | 9 | W | Exposed | 0.309 | 74.289 |

<sup>†</sup>RSA=Relative Surface Accessibility, <sup>‡</sup>ASA=Absolute Surface Accessibility

#### 3.2.6 Prediction of Change in Stability of the Mutated Protein by Consensus Prediction

The amino acid substitutions were next analyzed by protein stability prediction tool, iStable 2.0. iStable 2.0 predicted 47 substitutions as destabilizing (Table 3).

**Table 3:** Stability Prediction of the Mutated Proteins by iStable 2.0 and DynaMut.

| AAS | iStable 2.0 |  | DynaMut |  | Missense3D |  |
| --- | --- | --- | --- | --- | --- | --- |
| | Confidence Score | Stability | $\Delta\Delta G$ (kcal/mol) | Stability | Structural Damage | Comment |
| L58P | -1.7689416 | Decrease | -1.348 | Destabilizing | No |  |
| T72K | -1.0673743 | Decrease | 0.677 | Stabilizing | Yes | Buried charge introduced |
| T72M | -0.067792356 | Increase | 0.472 | Stabilizing | No |  |
| G80R | -1.4305459 | Decrease | 0.826 | Stabilizing | Yes | Clash alert, Buried charge introduced, Disallowed phi/psi alert, Buried Gly replaced |
| F84L | -1.3939497 | Decrease | -0.815 | Destabilizing | No |  |
| P85L | -0.40284556 | Increase | -0.257 | Destabilizing | Yes | Cis proline replaced |
| P85S | -1.9663911 | Decrease | -0.991 | Destabilizing | Yes | Cis proline replaced |
| V91A | -2.363688 | Decrease | -1.751 | Destabilizing | No |  |
| G93S | -1.005738 | Decrease | -0.948 | Destabilizing | No |  |
| G93D | -1.0917563 | Decrease | -0.592 | Destabilizing | No |  |
| G93V | -1.1831716 | Increase | -0.153 | Destabilizing | No |  |
| P96S | -1.0492245 | Decrease | 1.117 | Stabilizing | No |  |
| Y100C | -1.5898333 | Decrease | -0.83 | Destabilizing | No |  |
| Y100F | -1.0702753 | Decrease | 0.061 | Stabilizing | No |  |
| M104T | -3.143975 | Decrease | -0.444 | Destabilizing | No |  |
| D105N | -0.9833634 | Decrease | 0.443 | Stabilizing | Yes | Buried salt bridge breakage |
| V107E | -1.7875004 | Decrease | 0.555 | Stabilizing | No |  |
| P108T | -0.82126737 | Decrease | 0.924 | Stabilizing | No |  |
| D110E | -0.68696284 | Decrease | -1.513 | Destabilizing | No |  |
| D110N | -0.63278186 | Decrease | -1.912 | Destabilizing | No |  |
| D111Y | -0.11971706 | Decrease | -0.260 | Destabilizing | No |  |
| R113I | -0.0076336265 | Increase | -0.166 | Destabilizing | No |  |
| R113K | -1.2718449 | Decrease | -0.449 | Destabilizing | No |  |
| Y114H | -2.133617 | Decrease | -1.729 | Destabilizing | No |  |
| W121R | -1.7060409 | Decrease | -1.785 | Destabilizing | Yes | Cavity altered (expansion of the cavity volume by 76.464 Å<sup>3</sup>) |
| G125R | -0.8936368 | Decrease | 1.276 | Stabilizing | Yes | Disallowed phi/psi alert |
| G133C | -0.7795974 | Decrease | 0.599 | Stabilizing | No |  |
| L135R | -0.6822803 | Decrease | -0.049 | Destabilizing | No |  |
| V137A | -1.5604656 | Decrease | -0.738 | Destabilizing | No |  |
| D140E | -0.47431314 | Increase | 0.009 | Stabilizing | No |  |
| P142S | -1.0563802 | Decrease | 0.281 | Stabilizing | Yes | Cis proline replaced |
| A143T | -1.4433032 | Decrease | 0.885 | Stabilizing | No |  |
| G145W | -0.910699 | Increase | -0.37 | Destabilizing | Yes | Clash alert, Buried Gly replaced |
| V153D | -2.2433639 | Decrease | -1.831 | Destabilizing | Yes | Buried hydrophilic residue introduced, Buried charge introduced |
| L160F | -1.6011777 | Decrease | -0.659 | Destabilizing | No |  |
| H170L | 0.340119 | Increase | 0.057 | Stabilizing | No |  |
| N174D | -0.6325445 | Decrease | -0.051 | Destabilizing | No |  |
| V186G | -2.7898633 | Decrease | -0.109 | Destabilizing | No |  |
| V186M | -0.37593216 | Decrease | 0.347 | Stabilizing | No |  |
| D189Y | -1.0340729 | Decrease | -0.006 | Destabilizing | No |  |
| H204D | -1.2200044 | Decrease | -1.702 | Destabilizing | No |  |
| V205F | -1.4460453 | Decrease | 0.454 | Stabilizing | No |  |
| E208D | -1.1203609 | Decrease | -1.772 | Destabilizing | Yes | Buried H-bond breakage |
| A213T | -1.3542162 | Decrease | -1.086 | Destabilizing | No |  |
| I222N | -2.3990505 | Decrease | -1.566 | Destabilizing | Yes | Buried hydrophilic residue introduced |
| T223M | -0.053840578 | Decrease | 0.863 | Stabilizing | No |  |
| K226R | -0.55279315 | Decrease | -0.600 | Destabilizing | No |  |
| K226E | -1.2064422 | Decrease | -1.730 | Destabilizing | No |  |
| F232V | -0.04029095 | Decrease | -0.108 | Destabilizing | No |  |
| R237P | -1.6782372 | Decrease | -0.338 | Destabilizing | No |  |
| R237Q | -0.98735225 | Decrease | -0.627 | Destabilizing | No |  |
| R237W | -0.68915725 | Increase | 0.903 | Stabilizing | No |  |
| S261C | -0.1880107 | Increase | 1.311 | Stabilizing | Yes | Cavity altered (expansion of the cavity volume by 113.616 Å<sup>3</sup>) |
| C323Y | -0.45005268 | Increase | 0.91 | Stabilizing | Yes | Clash alert |
| P337H | -0.93664145 | Decrease | 0.614 | Stabilizing | No |  |
| R375W | -0.0087457895 | Increase | -0.659 | Destabilizing | No |  |
| W401C | -0.64045143 | Decrease | 1.108 | Stabilizing | Yes | Cavity altered (expansion of the cavity volume by 103.68 Å<sup>3</sup>) |

#### 3.2.7 Prediction of the Effects of Amino Acid Substitutions on the Structural Integrity of TBX5 Protein by Missense3D and Normal Mode Analysis

The amino acid substitutions were subsequently analyzed by Missense3D to see how these amino acid substitutions would affect protein tertiary structure. 15 substituted structures were predicted to be ‘Damaging’ by Missense3D (Table 3). Details of Missense3D predictions can be found in Supplementary Figures 3-17. DynaMut predicted 37 substitutions as destabilizing for the protein. Among the 15 substitutions predicted to be detrimental by Misssense3D, 13 were also predicted as damaging by either iStable2.0 or DynaMut, and 5 were predicted to be destabilizing by both (Figure 5). Table 3 provides predictions by these tools.

**Fig. 5:**
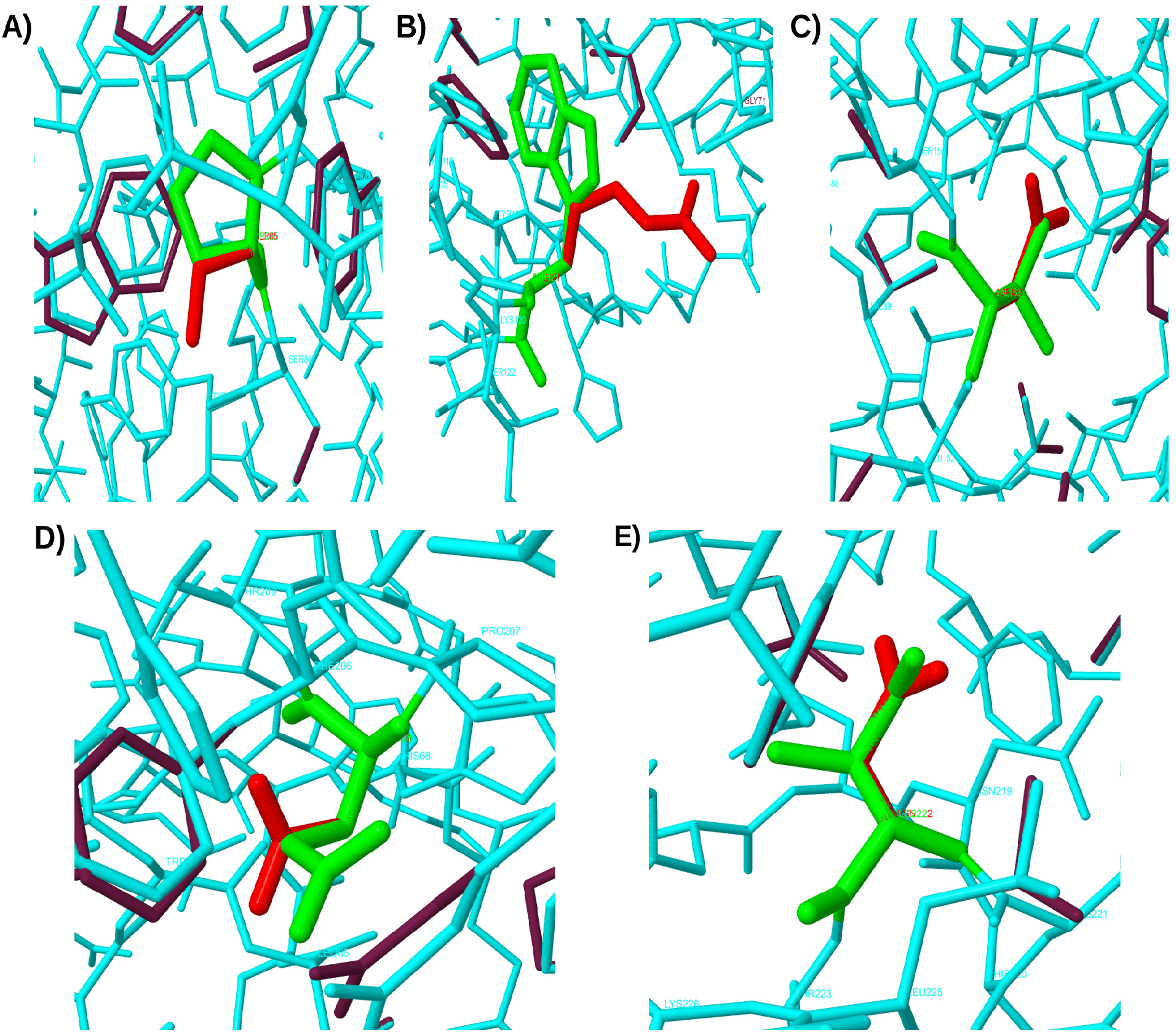
Close-up views of the 5 substitutions predicted to be damaging by iStable 2.0, Missense3D, and DynaMut. These 5 substitutions are A) P85S, B) W121R, C) V153D, D) E208D, and E) I222N. Native amino acids are shown in green and mutated amino acids are shown in red in this figure.

#### 3.2.8 Molecular Dynamics Simulation

Differences between the Cα RMSD profile of native structure (5BQD_A) and that of mutant structures (downloaded from Missense3D) are visible in Figure 6 (A). RMSD profiles of the mutant structures showed more deviations than the native structure. From the Cα RMSF profiles (Figure 6 (B)), it can be found that residue-wise RMSF values of the mutant structures were greater for all residues than the native structure (residue N192 is an exception). Comparing RMSF values of specific mutant residues with their native counterparts (Figure 6 (C)) it was found that W121R, V153D, and E208D showed considerably greater fluctuation. Fluctuation of P85S and I222N was close to their respective native residues. Greater deviations in the RMSD and RMSF profiles support unstable nature of the TBX5 mutant proteins and thus their likelihood to cause disease.

**Fig. 6:**
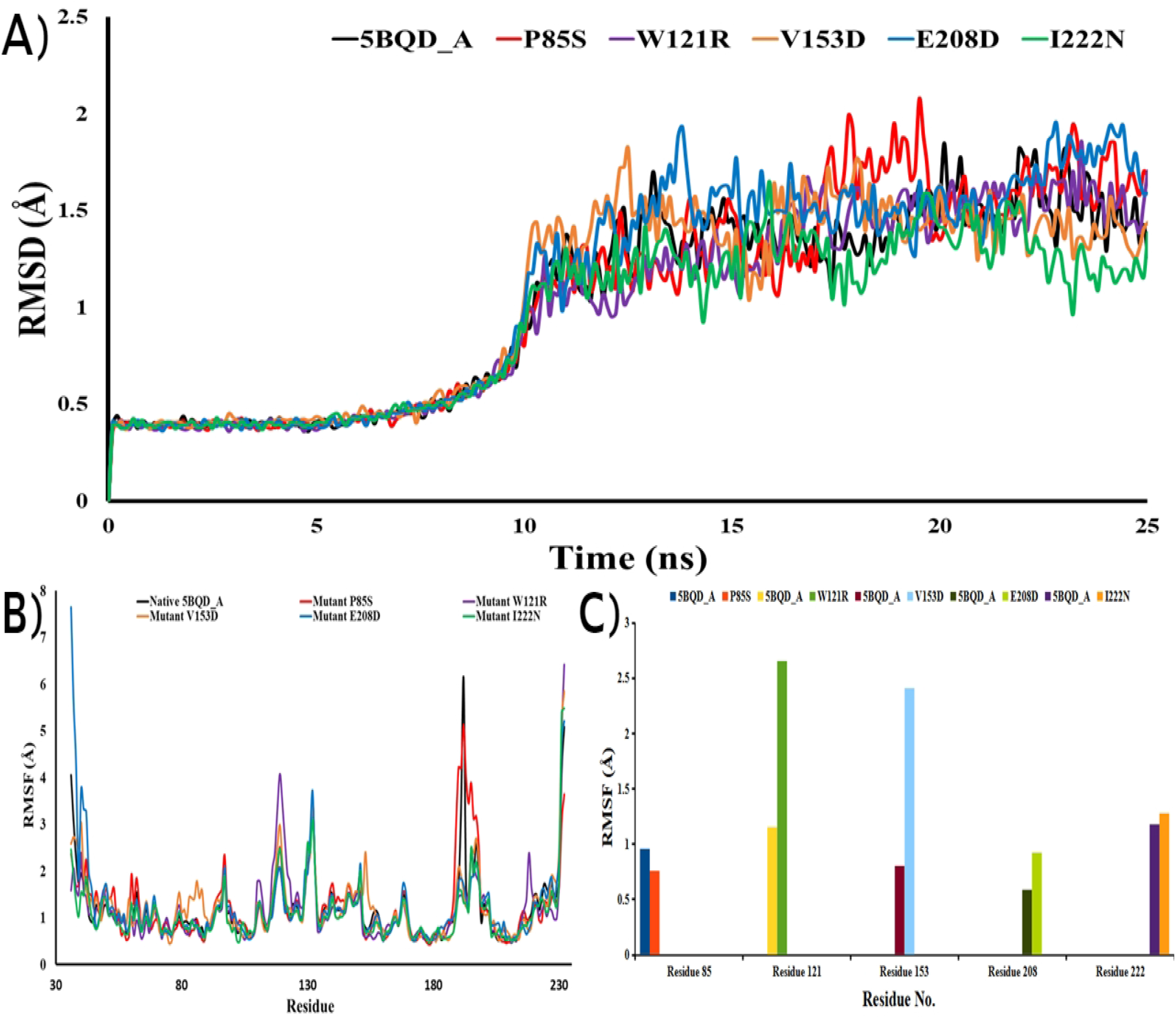
(A) Cα RMSD values of the native and mutant TBX5 proteins. (B) Residue-wise Cα RMSF values of native and mutant TBX5 proteins. (C) Comparison between the Cα RMSF values of native and mutant residues.

#### 3.2.8 Prediction of Post-translational Modification Site Missense SNPs by AWESOME

Among the 58 high-confidence deleterious SNPs, AWESOME predicted P108T, D111Y, and A143T substitutions could create new phosphorylation sites (AWESOME scores are -0.711, -1.458 and - 0.05, respectively), and S261C substitution can cause loss of a phosphorylation site (AWESOME score 2.752). R237P, R237Q, R237W, and R375W can cause loss of methylation sites (AWESOME scores are 0.834, 0.834, 0.834, and 0.768, respectively). S261C substitution can cause loss of serine mediated O-linked N-acetylgalactosamine (AWESOME score 0.8291) and O-linked N-acetylglucosamine formation (AWESOME score 0.4322). P108T and A143T substitutions can cause a gain of threonine mediated O-linked N-acetylglucosamine formation (AWESOME scores are -0.3507 and -0.354, respectively).

### 3.3 Evidence of Deleteriousness from *in vitro* Experiments

Among our identified 58 missense SNPs, G80R [76], G125R [77], T223M [78], R237P [79], R237Q, and R237W [76] have been confirmed experimentally to be responsible for HOS. G80R, R237P, R237Q, and R237W cause loss of function, whereas G125R leads to function [79].

### 3.4 Identification and Analyzing Nonsense and Stop Lost SNPs by PredictSNP2

2 stop gained (Nonsense), and 1 stop PredictSNP2 analyzed lost SNPs. It predicted the nonsense mutations as deleterious, and the stop lost mutation as neutral. Q156Ter nonsense mutation is significant because it disrupts the highly conserved T-box (amino acid position 58-238). Table 4 shows the results from PredictSNP2 server.

**Table 4:** Prediction about Nonsense and Stop Lost SNPs by PredictSNP2.

| Variant | PredictSNP2 |  | CADD |  | DANN |  | FATHMM |  | FunSeq2 |  | GWAVA |  |
| --- | --- | --- | --- | --- | --- | --- | --- | --- | --- | --- | --- | --- |
|  | P | S | P | S | P | S | P | S | P | S | P | S |
| W514Ter | Del | 0.6777 | Del | 46 | Del | 0.9959 | Del | 0.9623 | Del | 5 | ? | 0.39 |
| Q156Ter | Del | 1 | Del | 38 | Del | 0.9984 | Del | 0.9167 | Del | 5 | Del | 0.5 |
| Ter519Q | Neu | -0.1146 | Neu | 14.06 | Neu | 0.758 | Del | 0.9717 | Neu | 0 | Del | 0.31 |

P= Prediction, S= Score, Del= Deleterious & Neu= Neutral

### 3.5 Prediction of the effects of UTR SNPs by UTRdb & MirSNP

UTR SNPs that could affect transcriptional motifs were retrieved from the UTRdb server. Three SNPs in the 3′ UTR (rs28730760, rs28730761, and rs883079) were found to be present in the polyadenylation (poly-A) sites and hence, may result in pathogenicity. In addition, UTRdb output returned six transcriptional motif matches in the 5′ UTR and three transcriptional motif matches in the 3′ UTR. Table 5 provides analysis from the UTRdb server.

**Table 5:** Transcriptional Motifs Present in *TBX5* Untranslated Regions.

| Signal Name | UTR region | Match total | NCBI Reference Sequence | Genomic Position (NCBI36/hg18) |
| --- | --- | --- | --- | --- |
| Uorf (Upstream Open Reading Frame) | 5' | 1 | NM_181486.4 | Chr12:113328041-113328106 |
|  |  | 2 | NM_000192.3 | Chr12:113330378-113330629 |
|  |  |  |  | Chr12:113330136-113330216 |
|  |  | 2 | NM_080717.3 | Chr12:113330378-113330629<br>Chr12:113330136-113330216 |
| IRES (Internal Ribosome Entry Site) | 5' | 1 | NM_000192.3 | Chr12:113326087-113330055 |
| CPE (Cytoplasmic polyadenylation element) | 3' | 1 | NM_181486.4 | Chr12:113276118-113276332 |
| K-Box | 3' | 1 | NM_181486.5 | Chr12:113276848-113276855 |
| PAS (Polyadenylation Signal) | 3' | 1 | NM_181486.6 | Chr12:113276118-113276144 |

3′ UTR SNPs that might create or break a miRNA target site or enhance or decrease miRNA binding to mRNA were predicted by the MirSNP server. A total number of 86 SNPs in the 3′ untranslated region of the *TBX5* gene were predicted to alter miRNA-mRNA binding. Supplementary Table 3 shows the results from the MirSNP server.

### 3.6 Predicting the effects of SNPs located in splice sites by HSF

Human Splicing Finder (HSF) web tool was employed to predict the effects of SNPs located in 5′ and 3′ splice sites. A total number of 11 splice site SNPs were subjected to assessment. All were found to have ‘probably no impact on splicing’ by HSF. Table 6 shows the predictions obtained from the HSF.

**Table 6:** Prediction about Splice Site SNPs by HSF.

| rsIDs | Alleles | Predicted Signal | Interpretation |
| --- | --- | --- | --- |
| **Splice Acceptor Variant** |  |  |  |
| rs1555226330 | C>A | No significant splicing motif alteration detected | This mutation has probably no impact on splicing |
| rs1031873727 | G>A | ESS Site broken | Alteration of an intronic ESS site, Probably no impact on splicing |
| rs1031873727 | G>T | ESS Site broken | Alteration of an intronic ESS site, Probably no impact on splicing |
| rs891464399 | G>A | New ESE Site | Creation of an intronic ESE site, Probably no impact on splicing |
| rs958951320 | C>T | No significant splicing motif alteration detected | This mutation has probably no impact on splicing |
| rs181078973 | A>C | No significant splicing motif alteration detected | This mutation has probably no impact on splicing |
| rs181078973 | A>G | No significant splicing motif alteration detected | This mutation has probably no impact on splicing |
| rs374919751 | C>G | New ESE Site | Creation of an intronic ESE site, Probably no impact on splicing |
| rs374919751 | C>T | No significant splicing motif alteration detected | This mutation has probably no impact on splicing |
| rs1288475235 | C>A | No significant splicing motif alteration detected | This mutation has probably no impact on splicing |
| rs1015550731 | T>C | ESS Site broken | Alteration of an intronic ESS site, Probably no impact on splicing |

### 3.7 Prediction about Deep Functional Intronic SNPs

19 among all the *TBX5* intron variants were predicted to affect transcription factor binding sites by FuncPred. These 19 SNPs were further analyzed by RegulomeDB 2.0 server, and only two among these SNPs (rs12827969 and rs61931002) can be considered functionally important RegulomeDB 2.0 score. Table 7 summarizes results from SNP FuncPred and RegulomeDB.

**Table 7:** Predictions Regarding Deep Intronic SNPs.

| SNP FuncPred |  |  |  |  |  | RegulomeDB |  |
| --- | --- | --- | --- | --- | --- | --- | --- |
| rsID | Chromosome | Position | Allele | Regulatory Potential | Conservation | Rank | Probability |
| rs11067100 | 12 | 113328237 | G/T | 0.271604 | 0.998 | 4 | 0.60906 |
| rs11067101 | 12 | 113328586 | A/G | 0.186436 | 0 | 4 | 0.60906 |
| rs11613605 | 12 | 113326220 | C/G | 0.099303 | 0 | 4 | 0.60906 |
| rs11837917 | 12 | 113329990 | G/T | 0.096837 | 0 | 4 | 0.60906 |
| rs12319868 | 12 | 113329396 | G/T | 0.150806 | 0 | 4 | 0.60906 |
| rs12372585 | 12 | 113330591 | A/G | 0.225674 | 0.19 | 4 | 0.60906 |
| rs12423887 | 12 | 113327989 | C/T | 0.410483 | 0.025 | 4 | 0.60906 |
| rs1248046 | 12 | 113329889 | T/C | 0 | 0 | 4 | 0.60906 |
| rs12827969 | 12 | 113326517 | A/T | 0.181987 | 1 | 2b | 0.80975 |
| rs1522368 | 12 | 113329052 | A/G | 0.148659 | 0.997 | 3a | 0.6893 |
| rs1522369 | 12 | 113329156 | A/G | 0.076716 | 0 | 4 | 0.60906 |
| rs2113436 | 12 | 113326379 | G/A | 0.469676 | 0.006 | 4 | 0.60906 |
| rs35203448 | 12 | 113328816 | A/G | 0.113463 | 0.5 | 4 | 0.60906 |
| rs3782467 | 12 | 113327309 | C/G | 0.000319 | 0.001 | 4 | 0.60906 |
| rs4553413 | 12 | 113328827 | A/G | 0.0709 | 0.567 | 4 | 0.60906 |
| rs57820630 | 12 | 113330823 | C/T | NA | NA | 4 | 0.60906 |
| rs61931002 | 12 | 113326665 | A/G | NA | NA | 2b | 0.64343 |
| rs736560 | 12 | 113327637 | A/C | 0.393482 | 0 | 4 | 0.60906 |
| rs7957609 | 12 | 113331519 | A/G | 0.316415 | 0.022 | 4 | 0.60906 |

**RegulomeDB ranks:** 1a-1f: likely to affect binding and linked to the expression of a gene target, 2a-2c: likely to affect binding, 3a-3b: less likely to affect binding, 4,5,6: minimal binding evidence.

### 3.8 3D Structure prediction of Full TBX5 Protein

Due to the unavailability of a TBX5 full-length protein crystal structure, full-length 3D models of TBX5 were created employing three different protein structure prediction algorithms, which were later energy minimized and refined. Supplementary Figure 1 shows cartoon presentations of the energy minimized and refined models.

### 3.9 Evaluation of TBX5 3D Models

3D Model evaluation is indispensable for checking the credibility of the generated model. PROCHECK assessed the 3D models generated in the current study, Verify 3D, and SWISS-MODEL Structure Assessment Tool. Ramachandran plot is a reliable qualitative indicator of protein structure [70]. PROCHECK calculated Ramachandran plots for the created models. A model is considered reasonable by PROCHECK if >90% residues are present in the most favored regions of the Ramachandran plot. Only one model (generated by Robetta and later energy minimized and refined) was found to fulfill this criterion. It has 94.5% of residues in the most favored regions. Supplementary Figure 2 shows the Ramachandran plots calculated by PROCHECK. Verify 3D is another tool that approves a model if its 80% or more amino acid residues score ≥0.2 in the 3D/1D profile. However, none of the models could fulfill this requirement. MolProbity score and QMEAN Z-score for our models were obtained from the SWISS-MODEL Structure Assessment Tool. A low MolProbity score and a high QMEAN Z-score are preferred. A negative QMEAN Z-score indicates instability in a protein structure [71]. All of our models have a negative QMEAN Z-score. This disagreement can be explained by the fact that presently available *ab initio* protein modeling tools predict poorly for proteins having more than 120 residues [80], and for our protein models, 279 residues (5BQD-A chain extends from 1-239 residues and the full-length protein has 518 residues) do not have any template. However, among the models, the energy minimized and refined Robetta model has the lowest MolProbity score and the least negative QMEAN Z-score. Considering Ramachandran plot results, MolProbity scores, and QMEAN Z-scores, it can be decided that Robetta outperformed two other protein structure prediction tools used in this study. Supplementary Table 4 summarizes the results obtained from PROCHECK, Verify 3D, and SWISS-MODEL Structure Assessment Tool, and Supplementary Figure 2 shows the Ramachandran plots obtained from PROCHECK.

## 4. Discussion

Recently bioinformatics approaches have become popular for identifying different human gene SNPs and predicting their structural and functional consequences. In the past few years, damaging SNPs of *CCR6* gene [81], *SMPX* gene [82], human aldehyde oxidase gene [83], *PTEN* gene [84], folate pathway genes [85], human bone morphogenetic protein receptor type 1A gene [86], *SKP2* gene [87], human adiponectin receptor 2 gene [88], *MMP9* gene [89], etc. have been identified with the help of different computational biology tools. With the rapid development of genomics, the number of reported SNPs in different databases is continuously increasing for the last few years. However, determining the SNPs that result in diseases is challenging. Various pathogenicity prediction tools can narrow this number to a list of high-confidence deleterious deleterious SNPs [90]. *TBX5* gene is an essential regulator of cardiac and upper limb development, and mutations in this gene are responsible for Holt-Oram syndrome, especially missense mutations. It is estimated that 1 in 100,000 neonates are born with Holt-Oram syndrome, and male and female children are affected equally. Since HOS is an autosomal dominant condition, it is imperative that all individuals with HOS and their parents and siblings get genetic counseling [91]. In the past years, some mutations, including SNPs, have been reported in association with the *TBX5* gene. However, most of the SNPs of this gene have not been characterized yet. In this study we have tried to fill this gap.

In our current study, we have analyzed all currently available *TBX5* SNPs. Since results from a single bioinformatics tool may be inconclusive, we have appointed 9 highly reliable prediction tools for identifying deleterious missense SNPs. The 58 deleterious missense SNPs that were initially identified by concordance among PROVEAN, SIFT, and PolyPhen-2 results were also damaged by 6 other tools (PredictSNP1, PredictSNP2, MetaLR, MetaSVM, and REVEL) with few exceptions. Among these 58 missense SNPs, 5 missense SNPs (G80R, G125R, T223M, R237P, R237Q, and R237W) have been experimentally found to be responsible for HOS [76–79].

The conservation analysis found that 55 among the 58 missense SNPs cause amino acid substitution in highly conserved positions (ConSurf score 7-9). This further corroborates the importance of these SNPs. Furthermore, 8 of these 58 missense SNPs (P108T, D111Y, A143T, R237P, R237Q, R237W, S261C, and R375W) were found to have the ability to modify post-translational sites.

We have also analyzed the effects of these 58 SNPs on the stability of native TBX5 protein. 15 SNPs were predicted to cause structural damage by Missense3D. G80R, T72K, and V153D substitutions introduce buried charges. V153D and I22N mutations substitute buried hydrophobic residues with hydrophilic residues. G80R and G145W replace a buried glycine, P85L, P85S, and P142S replace a *cis* proline, D105N breaks a buried salt bridge, and E208D disrupts a side-chain / side-chain H-bond and a side-chain / main-chain H-bond. W121R, S261C, and W401C cause an expansion in the internal protein cavity and thus make protein structure unstable. G80R, G145W, and C323Y substitutions trigger clash alerts. These three substitutions lead to a MolProbity clash score ≥30, and the increase in clash score is >18 compared to the wild-type. G80R and G125R mutations induce disallowed phi/psi alerts. 5 of these 15 substitutions (P85S, W121R, V153D, E208D, and I222N) were predicted to destabilize native protein structure by the consensus of Missense3D, DynaMut, and iStable 2.0. These SNPs which have not yet been reported [rs1565941579 (P85S), rs1269970792 (W121R), rs772248871 (V153D), rs769113870 (E208D), and rs1318021626 (I222N)] are most likely to cause HOS.

We have also employed 6 other tools to identify deleterious non-coding SNPs on the *TBX5* gene. 86 SNPs in the 3′ UTR of the *TBX5* gene were identified that could affect miRNA-mRNA binding. No significant SNP was identified in the splice sites of *TBX5,* and 2 deep intronic SNPs were identified that could affect transcription factor binding and gene expression.

Due to the unavailability of a full-size experimental TBX5 protein structure, 3D models were also predicted employing I-TASSER, Phyre2, and Robetta. Later, the 3D structures were evaluated by PROCHECK, Verify 3D, and SWISS-MODEL Structure Assessment Tool. A model is considered good if >90% of residues are present in the most favored regions of the Ramachandran plot. In the case of our *TBX5* models, only Robetta generated model passed this requirement (94.5%). On the other hand, verify 3D results and QMEAN Z-scores indicate poor quality of the generated models. This can be explained by the inability of the available *ab initio* protein modeling tools to predict correct structure of proteins having more than 120 residues [80]. These template-less regions are responsible for the deviations of our models from an ideal one.

## 5. Conclusion

This extensive *in silico* study attempts to find putatively most damaging *TBX5* SNPs. Our obtained results can guide further wet-lab studies of *TBX5* gene-related diseases and may potentiate finding cures in the future.

## Abbreviations

dbSNP: Database of Single Nucleotide Polymorphisms
HOS: Holt-Oram Syndrome
I-TASSER: Iterative Threading ASSEmbly Refinement
INDEL: Insertion-Deletion
NCBI: National Center for Biotechnology Information
PDB: Protein Data Bank
PD: Probably Damaging
PoD: Possibly Damaging
PROVEAN: PROtein Variation Effect Analyzer
PolyPhen-2: Polymorphism Phenotyping version 2
PTM: Post-translational Modification
REVEL: Rare Exome Variant Ensemble Learner
rs ID: reference SNP ID
SIFT: Sorting Intolerant From Tolerant
SNP: Single Nucleotide Polymorphism
TBX5: T-box 5 gene
TBX5: T-box 5 transcription factor (protein)
TF: Transcription Factor
UTR: Untranslated Region
VEP: Variant Effect Predictor
YASARA: Yet Another Scientific Artificial Reality Application

## Declarations

### Ethics approval and consent to participate

Not applicable.

### Consent for publication

Not applicable.

### Availability of data and material

All data generated or analyzed during this study are included in this paper and the supplementary file.

## References

1. Papaioannou VE. The T-box gene family: emerging roles in development, stem cells and cancer. Development. 2014;141: 3819–3833. doi:10.1242/dev.104471

2. Papaioannou VE. T-box genes in development: From hydra to humans. International Review of Cytology. 2001;207: 1–70. doi:10.1016/S0074-7696(01)07002-4

3. Packham EA, Brook JD. T-box genes in human disorders. Human Molecular Genetics. 2003;12: 37R – 44. doi:10.1093/hmg/ddg077

4. Ghosh TK, Brook JD, Wilsdon A. T-Box Genes in Human Development and Disease. Current Topics in Developmental Biology. 2017;122: 383–415. doi:10.1016/bs.ctdb.2016.08.006

5. Ghosh TK, Packham EA, Bonser AJ, Robinson TE, Cross SJ, Brook JD. Characterization of the TBX5 binding site and analysis of mutations that cause Holt–Oram syndrome. Human Molecular Genetics. 2001;10: 1983–1994. doi:10.1093/hmg/10.18.1983

6. Hiroi Y, Kudoh S, Monzen K, Ikeda Y, Yazaki Y, Nagai R, et al. Tbx5 associates with Nkx2-5 and synergistically promotes cardiomyocyte differentiation. Nature Genetics. 2001;28: 276–280. doi:10.1038/90123

7. Steimle JD, Moskowitz IP. TBX5: A Key Regulator of Heart Development. Current Topics in Developmental Biology. 2017;122: 195–221. doi:10.1016/bs.ctdb.2016.08.008

8. Rallis C, Bruneau BG, Del Buono J, Seidman CE, Seidman JG, Nissim S, et al. Tbx5 is required for forelimb bud formation and continued outgrowth. Development. 2003;130: 2741–2751. doi:10.1242/dev.00473

9. Hasson P, Del Buono J, Logan MPO. Tbx5 is dispensable for forelimb outgrowth. Development. 2007;134: 85–92. doi:10.1242/dev.02622

10. Običan S, Maggio L. Holt-Oram Syndrome. Obstetric Imaging: Fetal Diagnosis and Care. 2018; 557–559.e1. doi:10.1016/B978-0-323-44548-1.00132-7

11. Boogerd CJJ, Dooijes D, Ilgun A, Mathijssen IB, Hordijk R, van de Laar IMBH, et al. Functional analysis of novel TBX5 T-box mutations associated with Holt–Oram syndrome. Cardiovascular Research. 2010;88: 130–139. doi:10.1093/cvr/cvq178

12. Vanlerberghe C, Jourdain A-S, Ghoumid J, Frenois F, Mezel A, Vaksmann G, et al. Holt-Oram syndrome: clinical and molecular description of 78 patients with TBX5 variants. European Journal of Human Genetics. 2019;27: 360–368. doi:10.1038/s41431-018-0303-3

13. Braulke I, Herzog S, Thies U, Zoll B. Holt-Oram syndrome in four half-siblings with unaffected parents: brief clinical report. Clinical Genetics. 1991;39: 241–244. doi:https://doi.org/10.1111/j.1399-0004.1991.tb03021.x

14. RESERVED IU-AR. Orphanet: Holt Oram syndrome. [cited 19 Apr 2021]. Available: https://www.orpha.net/consor/cgi-bin/OC_Exp.php?Expert=392&lng=EN

15. Barisic I, Boban L, Greenlees R, Garne E, Wellesley D, Calzolari E, et al. Holt Oram syndrome: a registry-based study in Europe. Orphanet Journal of Rare Diseases. 2014;9: 156. doi:10.1186/s13023-014-0156-y

16. Yi Li Q, Newbury-Ecob RA, Terrett JA, Wilson DI, Curtis ARJ, Ho Yi C, et al. Holt-Oram syndrome is caused by mutations in TBX5, a member of the Brachyury (T) gene family. Nature Genetics. 1997;15: 21–29. doi:10.1038/ng0197-21

17. Yang J, Hu D, Xia J, Yang Y, Ying B, Hu J, et al. Three novel TBX5 mutations in Chinese patients with Holt-Oram syndrome. Am J Med Genet. 2000;92: 237–240. doi:10.1002/(sici)1096-8628(20000605)92:4<237::aid-ajmg2>3.0.co;2-g

18. Borozdin W, Acosta AMB-F, Seemanova E, Leipoldt M, Bamshad MJ, Unger S, et al. Contiguous hemizygous deletion of TBX5, TBX3, and RBM19 resulting in a combined phenotype of Holt-Oram and ulnar-mammary syndromes. American Journal of Medical Genetics Part A. 2006;140A: 1880–1886. doi:https://doi.org/10.1002/ajmg.a.31340

19. Patel C, Silcock L, McMullan D, Brueton L, Cox H. TBX5 intragenic duplication: a family with an atypical Holt–Oram syndrome phenotype. European Journal of Human Genetics. 2012;20: 863– 869. doi:10.1038/ejhg.2012.16

20. Guo Q, Shen J, Liu Y, Pu T, Sun K, Chen S. Exome Sequencing Identifies a c.148-1G>C Mutation of TBX5 in a Holt-Oram Family with Unusual Genotype-Phenotype Correlations. CPB. 2015;37: 1066–1074. doi:10.1159/000430232

21. Ríos-Serna LJ, Díaz-Ordoñez L, Candelo E, Pachajoa H. A novel de novo *TBX5* mutation in a patient with Holt-Oram syndrome. Application of Clinical Genetics. 2018;11:157–162. doi:10.2147/TACG.S183418

22. Borozdin W, Acosta AMBF, Bamshad MJ, Botzenhart EM, Froster UG, Lemke J, et al. Expanding the spectrum of TBX5 mutations in Holt-Oram syndrome: detection of two intragenic deletions by quantitative real time PCR, and report of eight novel point mutations. Human Mutation. 2006;27: 975–976. doi:https://doi.org/10.1002/humu.9449

23. Alaaeldin G. Fayez, Nora N. Esmaiel, Arwa A. El Darsh, Engy A. Ashaat, Alaa K. Kamel, Ghada M. Shehata, et al. prediction of the functional consequences of a novel homozygous TBX5 variant in isolated AVSD patient. Journal of Innovations in Pharmaceutical and Biological Sciences. 2017;4: 97–105.

24. Zhang X-L, Qiu X-B, Yuan F, Wang J, Zhao C-M, Li R-G, Xu L, Xu Y-J, Shi H-Y, Hou X-M, Qu X-K, Xu Y-W, Yang Y-Q. TBX5 loss-of-function mutation contributes to familial dilated cardiomyopathy. Biochemical and Biophysical Research Communications. 2015;459: 166–171. doi:10.1016/j.bbrc.2015.02.094

25. Zhou W, Zhao L, Jiang J-Q, Jiang W-F, Yang Y-Q, Qiu X-B. A novel TBX5 loss-of-function mutation associated with sporadic dilated cardiomyopathy. International Journal of Molecular Medicine. 2015;36: 282–288. doi:10.3892/ijmm.2015.2206

26. Zhang R, Tian X, Gao L, Li H, Yin X, Dong Y, et al. Common Variants in the TBX5 Gene Associated with Atrial Fibrillation in a Chinese Han Population. PLOS ONE. 2016;11: e0160467. doi:10.1371/journal.pone.0160467

27. Baban A, Postma AV, Marini M, Trocchio G, Santilli A, Pelegrini M, et al. Identification of TBX5 mutations in a series of 94 patients with Tetralogy of Fallot. American Journal of Medical Genetics Part A. 2014;164: 3100–3107. doi:https://doi.org/10.1002/ajmg.a.36783

28. Wang F, Liu D, Zhang R-R, Yu L-W, Zhao J-Y, Yang X-Y, et al. A TBX5 3′UTR variant increases the risk of congenital heart disease in the Han Chinese population. Cell Discov. 2017;3: 17026. doi:10.1038/celldisc.2017.26

29. Choi Y, Sims GE, Murphy S, Miller JR, Chan AP. Predicting the Functional Effect of Amino Acid Substitutions and Indels. PLOS ONE. 2012;7: e46688. doi:10.1371/journal.pone.0046688

30. Ng PC, Henikoff S. SIFT: predicting amino acid changes that affect protein function. Nucleic Acids Research. 2003;31: 3812–3814. doi:10.1093/nar/gkg509

31. Kumar P, Henikoff S, Ng PC. Predicting the effects of coding non-synonymous variants on protein function using the SIFT algorithm. Nature Protocols. 2009;4: 1073–1081. doi:10.1038/nprot.2009.86

32. Adzhubei IA, Schmidt S, Peshkin L, Ramensky VE, Gerasimova A, Bork P, et al. A method and server for predicting damaging missense mutations. Nature Methods. 2010;7: 248–249. doi:10.1038/nmeth0410-248

33. Pejaver V, Urresti J, Lugo-Martinez J, Pagel KA, Lin GN, Nam H-J, et al. Inferring the molecular and phenotypic impact of amino acid variants with MutPred2. Nature Communications. 2020;11: 5918. doi:10.1038/s41467-020-19669-x

34. Bendl J, Musil M, Štourač J, Zendulka J, Damborský J, Brezovský J. PredictSNP2: A Unified Platform for Accurately Evaluating SNP Effects by Exploiting the Different Characteristics of Variants in Distinct Genomic Regions. PLOS Computational Biology. 2016;12: e1004962. doi:10.1371/journal.pcbi.1004962

35. Bendl J, Stourac J, Salanda O, Pavelka A, Wieben ED, Zendulka J, et al. PredictSNP: Robust and Accurate Consensus Classifier for Prediction of Disease-Related Mutations. PLOS Computational Biology. 2014;10: e1003440. doi:10.1371/journal.pcbi.1003440

36. Dong C, Wei P, Jian X, Gibbs R, Boerwinkle E, Wang K, et al. Comparison and integration of deleteriousness prediction methods for non-synonymous SNVs in whole exome sequencing studies. Human Molecular Genetics. 2015;24: 2125–2137. doi:10.1093/hmg/ddu733

37. Ioannidis NM, Rothstein JH, Pejaver V, Middha S, McDonnell SK, Baheti S, et al. REVEL: An Ensemble Method for Predicting the Pathogenicity of Rare Missense Variants. The American Journal of Human Genetics. 2016;99: 877–885. doi:10.1016/j.ajhg.2016.08.016

38. Ashkenazy H, Abadi S, Martz E, Chay O, Mayrose I, Pupko T, et al. ConSurf 2016: an improved methodology to estimate and visualize evolutionary conservation in macromolecules. Nucleic Acids Res. 2016;44: W344–W350. doi:10.1093/nar/gkw408

39. Jubb HC, Pandurangan AP, Turner MA, Ochoa-Montaño B, Blundell TL, Ascher DB. Mutations at protein-protein interfaces: Small changes over large surfaces have large impacts on human health. Progress in Biophysics and Molecular Biology. 2017;128: 3–13. doi:10.1016/j.pbiomolbio.2016.10.002

40. van Wijk R, Rijksen G, Huizinga EG, Nieuwenhuis HK, van Solinge WW. HK Utrecht: missense mutation in the active site of human hexokinase associated with hexokinase deficiency and severe nonspherocytic hemolytic anemia. Blood. 2003;101: 345–347. doi:10.1182/blood-2002-06-1851

41. Klausen MS, Jespersen MC, Nielsen H, Jensen KK, Jurtz VI, Sønderby CK, et al. NetSurfP-2.0: Improved prediction of protein structural features by integrated deep learning. Proteins: Structure, Function, and Bioinformatics. 2019;87: 520–527. doi:https://doi.org/10.1002/prot.25674

42. Ittisoponpisan S, Islam SA, Khanna T, Alhuzimi E, David A, Sternberg MJE. Can Predicted Protein 3D Structures Provide Reliable Insights into whether Missense Variants Are Disease Associated? Journal of Molecular Biology. 2019;431: 2197–2212. doi:10.1016/j.jmb.2019.04.009

43. Stefl S, Nishi H, Petukh M, Panchenko AR, Alexov E. Molecular Mechanisms of Disease-Causing Missense Mutations. Journal of Molecular Biology. 2013;425: 3919–3936. doi:10.1016/j.jmb.2013.07.014

44. Chen C-W, Lin M-H, Liao C-C, Chang H-P, Chu Y-W. iStable 2.0: Predicting protein thermal stability changes by integrating various characteristic modules. Computational and Structural Biotechnology Journal. 2020;18: 622–630. doi:10.1016/j.csbj.2020.02.021

45. Rodrigues CH, Pires DE, Ascher DB. DynaMut: predicting the impact of mutations on protein conformation, flexibility and stability. Nucleic Acids Research. 2018;46: W350–W355. doi:10.1093/nar/gky300

46. Doerr S, Harvey MJ, Noé F, De Fabritiis G. HTMD: High-Throughput Molecular Dynamics for Molecular Discovery. J Chem Theory Comput. 2016;12: 1845–1852. doi:10.1021/acs.jctc.6b00049

47. Yang Y, Peng X, Ying P, Tian J, Li J, Ke J, et al. AWESOME: a database of SNPs that affect protein post-translational modifications. Nucleic Acids Research. 2019;47: D874–D880. doi:10.1093/nar/gky821

48. Mignone F, Gissi C, Liuni S, Pesole G. [No title found]. Genome Biol. 2002;3: reviews0004.1. doi:10.1186/gb-2002-3-3-reviews0004

49. Steri M, Idda ML, Whalen MB, Orrù V. Genetic variants in mRNA untranslated regions. WIREs RNA. 2018;9: e1474. doi:10.1002/wrna.1474

50. Mohan RA, van Engelen K, Stefanovic S, Barnett P, Ilgun A, Baars MJH, et al. A mutation in the Kozak sequence of *GATA4* hampers translation in a family with atrial septal defects. Am J Med Genet. 2014;164: 2732–2738. doi:10.1002/ajmg.a.36703

51. von Bohlen AE, Böhm J, Pop R, Johnson DS, Tolmie J, Stücker R, et al. A mutation creating an upstream initiation codon in the *SOX9* 5′ UTR causes acampomelic campomelic dysplasia. Mol Genet Genomic Med. 2017;5: 261–268. doi:10.1002/mgg3.282

52. Witt H, Luck W, Hennies HC, Claßen M, Kage A, Laß U, et al. Mutations in the gene encoding the serine protease inhibitor, Kazal type 1 are associated with chronic pancreatitis. Nat Genet. 2000;25: 213–216. doi:10.1038/76088

53. Wen Y, Liu Y, Xu Y, Zhao Y, Hua R, Wang K, et al. Loss-of-function mutations of an inhibitory upstream ORF in the human hairless transcript cause Marie Unna hereditary hypotrichosis. Nature Genetics. 2009;41: 228–233. doi:10.1038/ng.276

54. Hudder A, Werner R. Analysis of a Charcot-Marie-Tooth Disease Mutation Reveals an Essential Internal Ribosome Entry Site Element in the Connexin-32 Gene. Journal of Biological Chemistry. 2000;275: 34586–34591. doi:10.1074/jbc.M005199200

55. Nicolas G, Wallon D, Goupil C, Richard A-C, Pottier C, Dorval V, et al. Mutation in the 3’untranslated region of APP as a genetic determinant of cerebral amyloid angiopathy. Eur J Hum Genet. 2016;24: 92–98. doi:10.1038/ejhg.2015.61

56. Grillo G, Turi A, Licciulli F, Mignone F, Liuni S, Banfi S, et al. UTRdb and UTRsite (RELEASE 2010): a collection of sequences and regulatory motifs of the untranslated regions of eukaryotic mRNAs. Nucleic Acids Research. 2010;38: D75–D80. doi:10.1093/nar/gkp902

57. Bartel DP. MicroRNAs: Genomics, Biogenesis, Mechanism, and Function. Cell. 2004;116: 281–297. doi:10.1016/S0092-8674(04)00045-5

58. Moszyńska A, Gebert M, Collawn JF, Bartoszewski R. SNPs in microRNA target sites and their potential role in human disease. Open Biology. 7: 170019. doi:10.1098/rsob.170019

59. Liu C, Zhang F, Li T, Lu M, Wang L, Yue W, et al. MirSNP, a database of polymorphisms altering miRNA target sites, identifies miRNA-related SNPs in GWAS SNPs and eQTLs. BMC Genomics. 2012;13: 661. doi:10.1186/1471-2164-13-661

60. Anna A, Monika G. Splicing mutations in human genetic disorders: examples, detection, and confirmation. J Appl Genetics. 2018;59: 253–268. doi:10.1007/s13353-018-0444-7

61. Wang G-S, Cooper TA. Splicing in disease: disruption of the splicing code and the decoding machinery. Nature Reviews Genetics. 2007;8: 749–761. doi:10.1038/nrg2164

62. Desmet F-O, Hamroun D, Lalande M, Collod-Béroud G, Claustres M, Béroud C. Human Splicing Finder: an online bioinformatics tool to predict splicing signals. Nucleic Acids Research. 2009;37: e67–e67. doi:10.1093/nar/gkp215

63. Vaz-Drago R, Custódio N, Carmo-Fonseca M. Deep intronic mutations and human disease. Hum Genet. 2017;136: 1093–1111. doi:10.1007/s00439-017-1809-4

64. Xu Z, Taylor JA. SNPinfo: integrating GWAS and candidate gene information into functional SNP selection for genetic association studies. Nucleic Acids Research. 2009;37: W600–W605. doi:10.1093/nar/gkp290

65. Boyle AP, Hong EL, Hariharan M, Cheng Y, Schaub MA, Kasowski M, et al. Annotation of functional variation in personal genomes using RegulomeDB. Genome Research. 2012;22: 1790– 1797. doi:10.1101/gr.137323.112

66. Yang J, Zhang Y. I-TASSER server: new development for protein structure and function predictions. Nucleic Acids Res. 2015;43: W174–W181. doi:10.1093/nar/gkv342

67. Kelley LA, Mezulis S, Yates CM, Wass MN, Sternberg MJE. The Phyre2 web portal for protein modeling, prediction and analysis. Nat Protoc. 2015;10: 845–858. doi:10.1038/nprot.2015.053

68. Song Y, DiMaio F, Wang RY-R, Kim D, Miles C, Brunette T, et al. High-Resolution Comparative Modeling with RosettaCM. Structure. 2013;21: 1735–1742. doi:10.1016/j.str.2013.08.005

69. Krieger E, Joo K, Lee J, Lee J, Raman S, Thompson J, et al. Improving physical realism, stereochemistry, and side-chain accuracy in homology modeling: Four approaches that performed well in CASP8: High-Resolution Homology Modeling. Proteins. 2009;77: 114–122. doi:10.1002/prot.22570

70. Laskowski RA, MacArthur MW, Thornton JM. PROCHECK : validation of protein-structure coordinates. 2012. pp. 684–687. doi:10.1107/97809553602060000882

71. Lüthy R, Bowie JU, Eisenberg D. Assessment of protein models with three-dimensional profiles. Nature. 1992;356: 83–85. doi:10.1038/356083a0

72. Williams CJ, Headd JJ, Moriarty NW, Prisant MG, Videau LL, Deis LN, et al. MolProbity: More and better reference data for improved all-atom structure validation. Protein Science. 2018;27: 293–315. doi:https://doi.org/10.1002/pro.3330

73. Studer G, Rempfer C, Waterhouse AM, Gumienny R, Haas J, Schwede T. QMEANDisCo— distance constraints applied on model quality estimation. Elofsson A, editor. Bioinformatics. 2020;36: 1765–1771. doi:10.1093/bioinformatics/btz828

74. Benkert P, Biasini M, Schwede T. Toward the estimation of the absolute quality of individual protein structure models. Bioinformatics. 2011;27: 343–350. doi:10.1093/bioinformatics/btq662

75. Clustered RefSNPs (rs) and Other Data Computed in House. National Center for Biotechnology Information (US); 2005. Available: https://www.ncbi.nlm.nih.gov/books/NBK44417/

76. Basson CT, Huang T, Lin RC, Bachinsky DR, Weremowicz S, Vaglio A, et al. Different TBX5 interactions in heart and limb defined by Holt-Oram syndrome mutations. Proceedings of the National Academy of Sciences. 1999;96: 2919–2924. doi:10.1073/pnas.96.6.2919

77. Postma Alex V., van de Meerakker Judith B.A., Mathijssen Inge B., Barnett Phil, Christoffels Vincent M., Ilgun Aho, et al. A Gain-of-Function TBX5 Mutation Is Associated With Atypical Holt–Oram Syndrome and Paroxysmal Atrial Fibrillation. Circulation Research. 2008;102: 1433– 1442. doi:10.1161/CIRCRESAHA.107.168294

78. Heinritz W. Identification of new mutations in the TBX5 gene in patients with Holt-Oram syndrome. Heart. 2005;91: 383–384. doi:10.1136/hrt.2004.036855

79. Boogerd CJJ, Dooijes D, Ilgun A, Mathijssen IB, Hordijk R, van de Laar IMBH, et al. Functional analysis of novel TBX5 T-box mutations associated with Holt-Oram syndrome. Cardiovascular Research. 2010;88: 130–139. doi:10.1093/cvr/cvq178

80. Lee J, Freddolino PL, Zhang Y. Ab Initio Protein Structure Prediction. In: J. Rigden D, editor. From Protein Structure to Function with Bioinformatics. Dordrecht: Springer Netherlands; 2017. pp. 3–35. doi:10.1007/978-94-024-1069-3_1

81. Akhtar M, Jamal T, Jamal H, Din JU, Jamal M, Arif M, et al. Identification of most damaging nsSNPs in human *CCR6* gene: In silico analyses. Int J Immunogenet. 2019;46: 459–471. doi:10.1111/iji.12449

82. Arifuzzaman Md, Mitra S, Das R, Hamza A, Absar N, Dash R. In silico analysis of non-synonymous single-nucleotide polymorphisms (nsSNPs) of the *SMPX* gene. Annals of Human Genetics. 2020;84: 54–71. doi:10.1111/ahg.12350

83. Coelho C, Muthukumaran J, Santos-Silva T, João Romão M. Systematic exploration of predicted destabilizing non-synonymous single nucleotide polymorphisms (nsSNPs) of human aldehyde oxidase: A Bio-informatics study. Pharmacol Res Perspect. 2019;7. doi:10.1002/prp2.538

84. Khan I, Ansari IA, Singh P, Dass J FP. Prediction of functionally significant single nucleotide polymorphisms in *PTEN* tumor suppressor gene: An *in silico* approach: Prediction in *PTEN* Tumor Suppressor Gene. Biotechnology and Applied Biochemistry. 2017;64: 657–666. doi:10.1002/bab.1483

85. Vohra M, Sharma AR, Paul B, Bhat MK, Satyamoorthy K, Rai PS. *In silico* characterization of functional single nucleotide polymorphisms of folate pathway genes. Annals of Human Genetics. 2018;82: 186–199. doi:10.1111/ahg.12231

86. Islam MdJ, Parves MdR, Mahmud S, Tithi FA, Reza MdA. Assessment of structurally and functionally high-risk nsSNPs impacts on human bone morphogenetic protein receptor type IA (BMPR1A) by computational approach. Computational Biology and Chemistry. 2019;80: 31–45. doi:10.1016/j.compbiolchem.2019.03.004

87. Hosen SMZ, Dash R, Junaid Md, Mitra S, Absar N. Identification and structural characterization of deleterious non-synonymous single nucleotide polymorphisms in the human SKP2 gene. Computational Biology and Chemistry. 2019;79: 127–136. doi:10.1016/j.compbiolchem.2019.02.003

88. Solayman Md, Saleh MdA, Paul S, Khalil MdI, Gan SH. In silico analysis of nonsynonymous single nucleotide polymorphisms of the human adiponectin receptor 2 (ADIPOR2) gene. Computational Biology and Chemistry. 2017;68: 175–185. doi:10.1016/j.compbiolchem.2017.03.005

89. Bhatnager R, Bhasin M, Dang AS. Comprehensive analysis of damage associated SNPs of MMP9 gene: A computational approach. Computational Biology and Chemistry. 2018;77: 97–108. doi:10.1016/j.compbiolchem.2018.09.008

90. P. S, D. TK, C. GPD, R. S, Zayed H. Determining the role of missense mutations in the POU domain of HNF1A that reduce the DNA-binding affinity: A computational approach. Verma C, editor. PLoS ONE. 2017;12: e0174953. doi:10.1371/journal.pone.0174953

91. Krauser AF, Ponnarasu S, Schury MP. Holt Oram Syndrome. StatPearls. Treasure Island (FL): StatPearls Publishing; 2021. Available: http://www.ncbi.nlm.nih.gov/books/NBK513339/

