## Supplementary for "Identification of deleterious single nucleotide polymorphism (SNP)s on the human *TBX5* gene & prediction of their structural & functional consequences: An *in silico* approach"

**for**

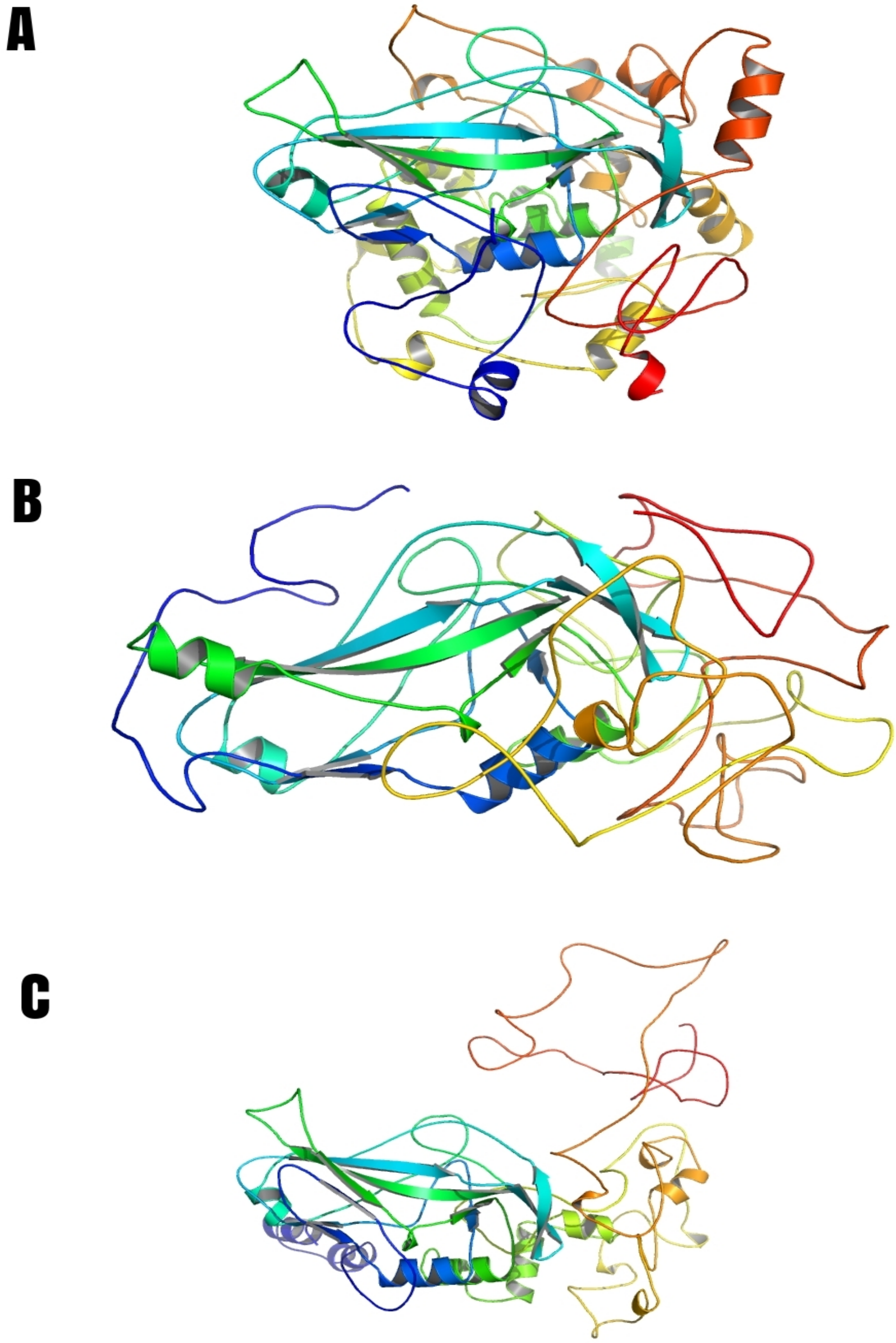

**Supplementary Figure 1:** Whole TBX5 model predicted by A) I-TASSER, B) Phyre2, and C) Robetta (Energy Minimized and Refined).

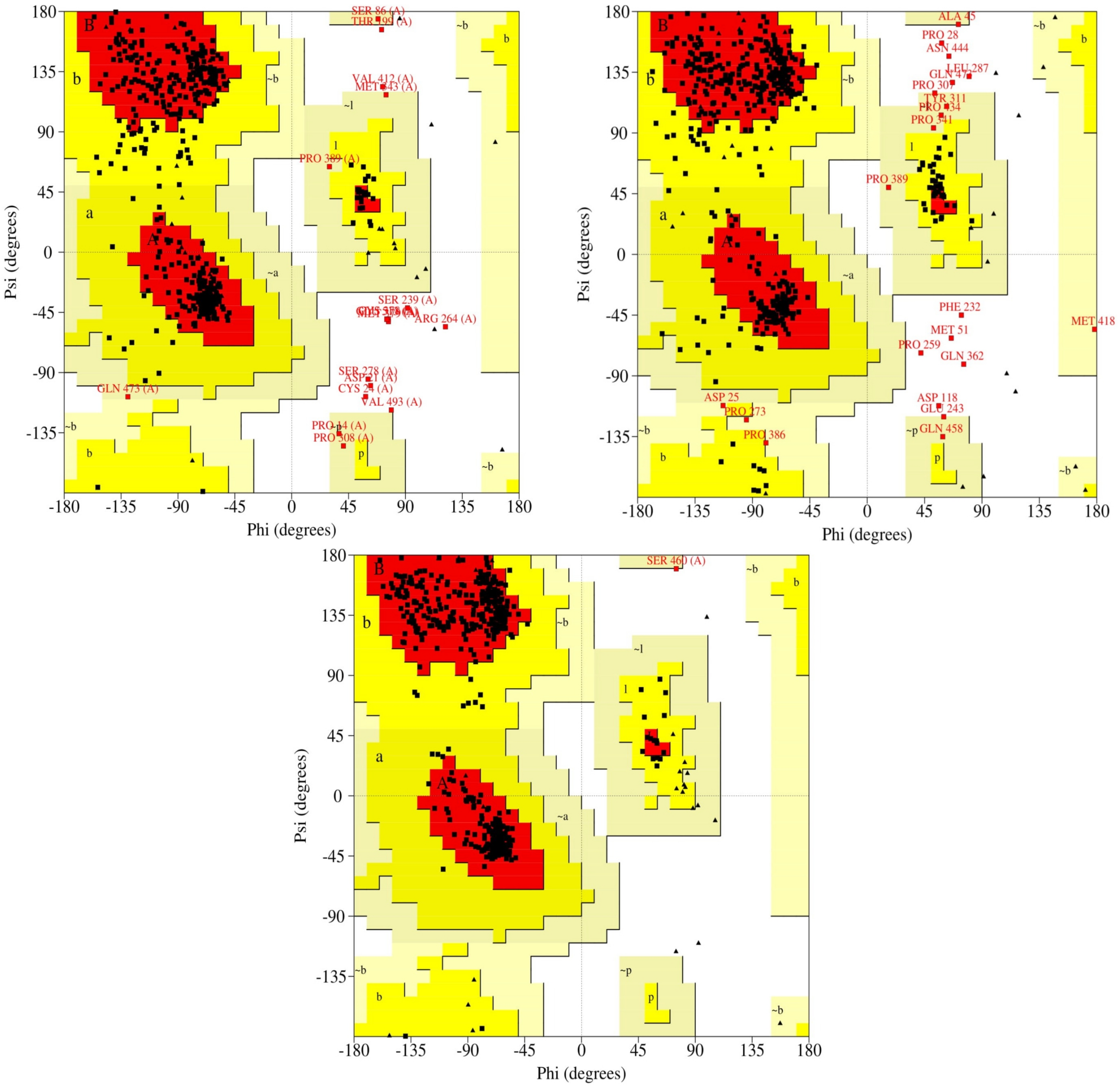

**Supplementary Figure 2:** A, B, and C are Ramachandran plots generated by PROCHECK of energy minimized and refined models produced by I-TASSER, Phyre2, and Robetta, respectively. Here, Red= most favored regions, Yellow= additional allowed regions, Light Yellow= generously allowed regions, and White= disallowed regions.

**Supplementary Figures 3-17** are close-up views of the 15 damaging mutations predicted by Missense3D. The wild-type chain color is cyan and the mutant chain color is rosewood. The wild-type residues are colored green and the mutant residues are colored red.

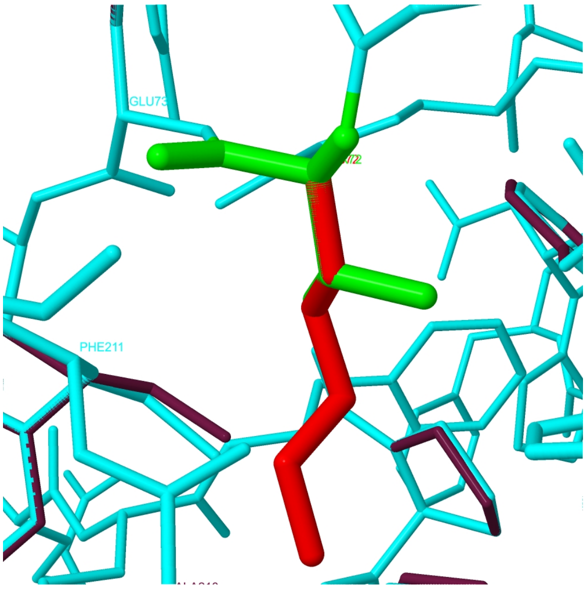

**Supplementary Figure 3:** Side chain replacement in T72K. This substitution replaces a buried uncharged residue (THR, RSA 3.5%) with a charged residue (LYS).

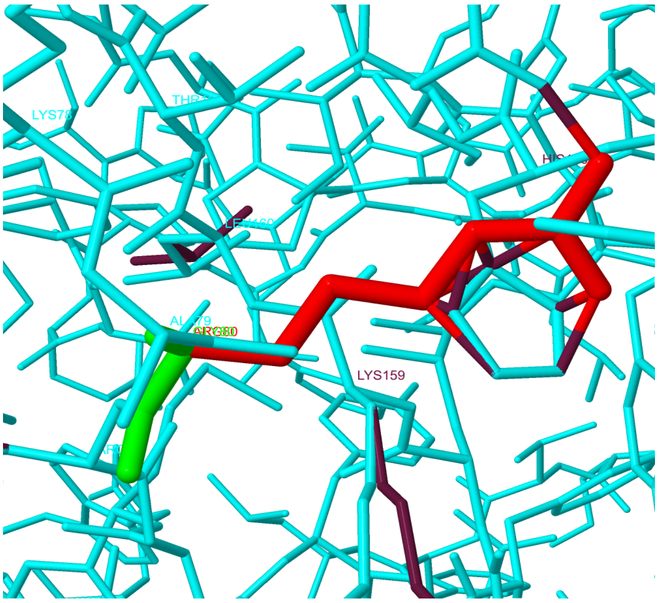

**Supplementary Figure 4:** Side chain replacement in G80R. This substitution triggers a clash alert. The local clash score for the wild-type residue is 19.29 and the local clash score for the mutant residue is 44.21. The mutant structure has a MolProbity clash score ≥30 and the increase in clash score is >18 compared to the wild-type. This substitution replaces a buried GLY residue (RSA 5.9%) with a buried ARG residue (RSA 4.8%). This substitution replaces a buried uncharged residue (GLY, RSA 5.9%) with a charged residue (ARG). This substitution also triggers disallowed phi/psi alert. The phi/psi angles are in the favored region of Ramachandran plot for the wild-type residue but outlier region for the mutant residue.

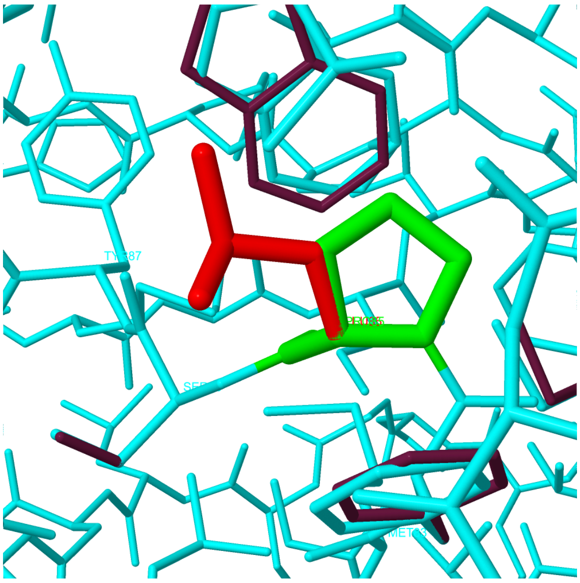

**Supplementary Figure 5:** Side chain replacement in P85L. This substitution replaces a wild-type *cis* proline.

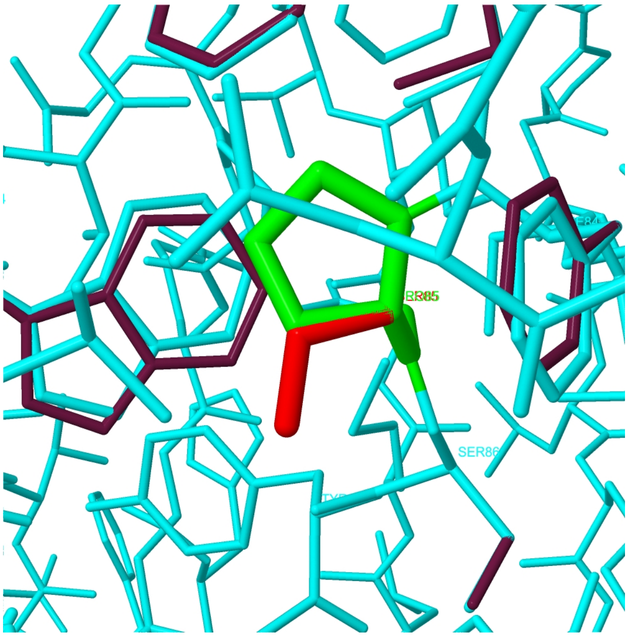

**Supplementary Figure 6:** Side chain replacement in P85S. This substitution replaces a wild-type *cis* proline.

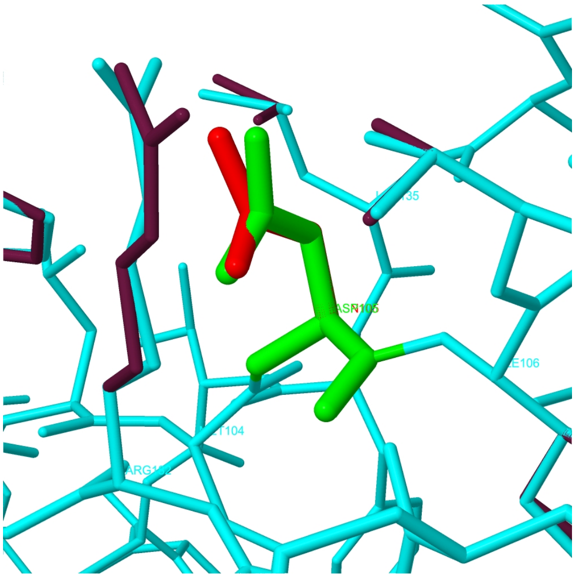

**Supplementary Figure 7:** Side chain replacement in D105N. This substitution disrupts a salt bridge formed between the OD1 atom of ASP 105 and the NE atom of ARG 182 (distance: 2.900 Å). The wild-type residue has an RSA of 7.3%.

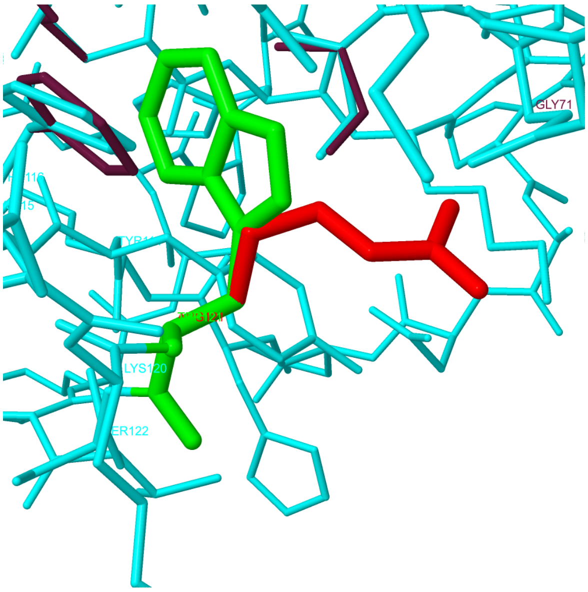

**Supplementary Figure 8:** Side chain replacement in W121R. This substitution leads to an expansion of the cavity volume by 76.464 Å^3.

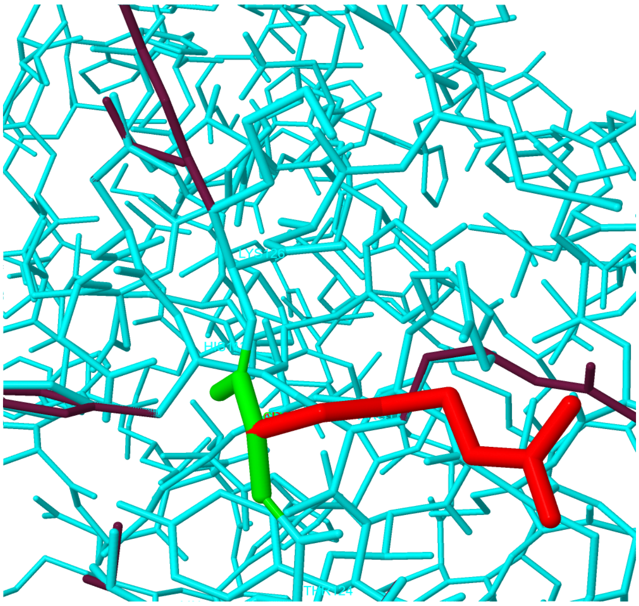

**Supplementary Figure 9:** Side chain replacement in G125R. This substitution triggers disallowed phi/psi alert. The phi/psi angles are in the favored region of Ramachandran plot for the wild-type residue but outlier region for the mutant residue.

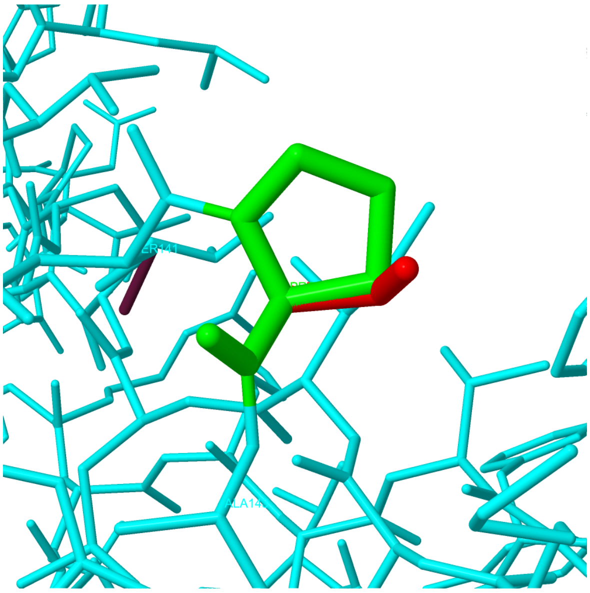

**Supplementary Figure 10:** Side chain replacement in P142S. This substitution replaces a wild-type *cis* proline.

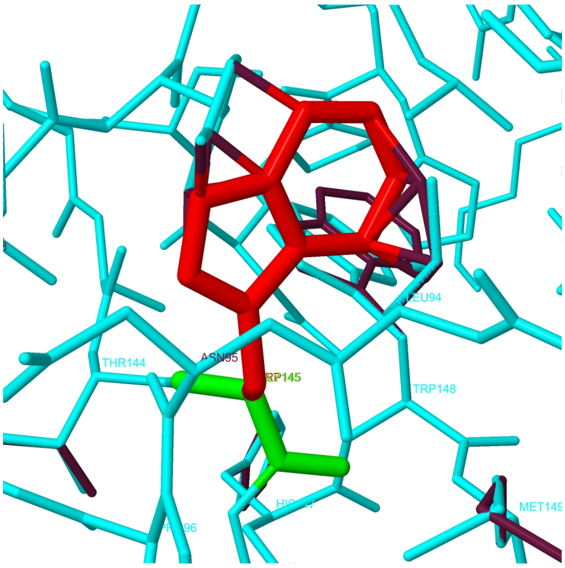

**Supplementary Figure 11:** Side chain replacement in G145W. This substitution triggers a clash alert. The local clash score for the wild-type residue is 26.68 and the local clash score for the mutant residue is 46.65. The mutant structure has a MolProbity clash score ≥30 and the increase in clash score is >18 compared to the wild-type. This substitution also replaces a buried GLY residue (RSA 0.0%) with a buried ARG residue (RSA 0.0%).

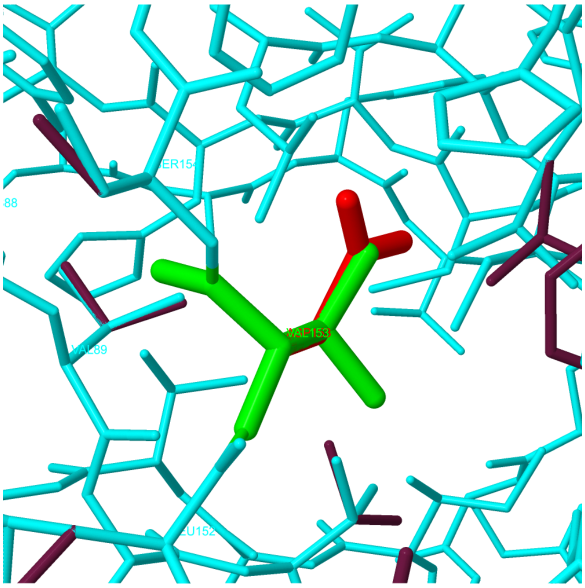

**Supplementary Figure 12:** Side chain replacement in V153D. This substitution replaces a buried hydrophobic residue (VAL, RSA 2.1%) with a hydrophilic residue (ASP, RSA 2.4%). This substitution also replaces a buried uncharged residue (VAL, RSA 2.1%) with a charged residue (ASP).

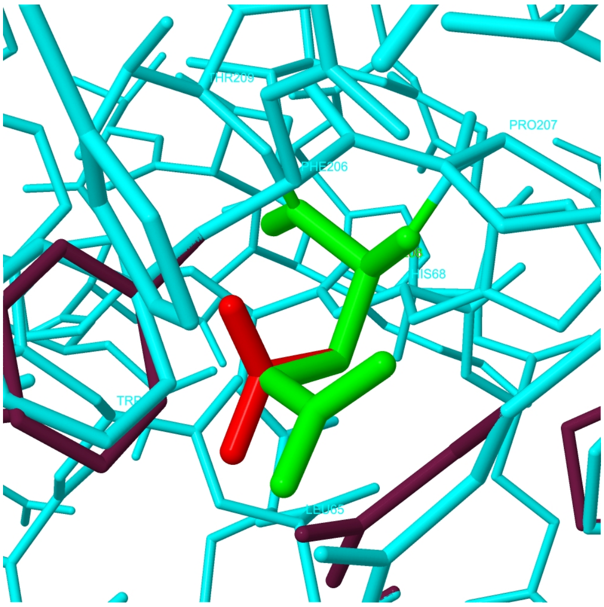

**Supplementary Figure 13:** Side chain replacement in E208D. GLU 208 is a buried residue (RSA 0.0%). This substitution disrupts a side-chain / side-chain H-bond (formed between the $NH_2$ group of ARG 61 and the OE2 atom of GLU 208; distance: 3.37 Å) and a side-chain / main-chain H-bond (formed between an N atom of GLU 208 and the OE1 atom of GLU 208; distance: 2.46 Å).

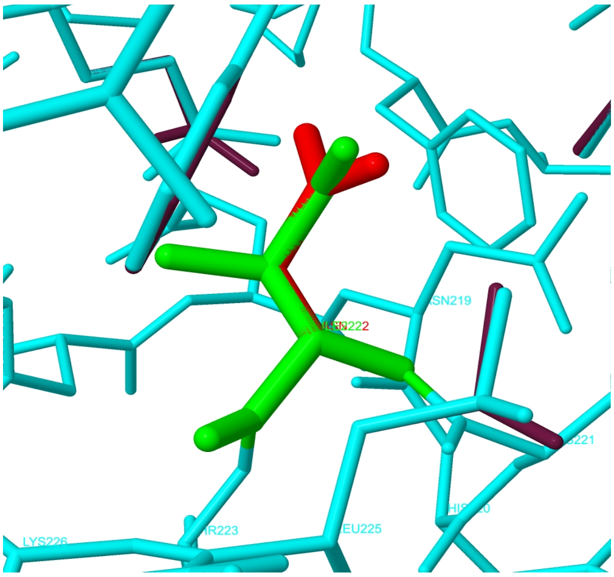

**Supplementary Figure 14:** Side chain replacement in I222N. This substitution replaces a buried hydrophobic residue (ILE, RSA 0.0%) with a hydrophilic residue (ASN, RSA 0.0%).

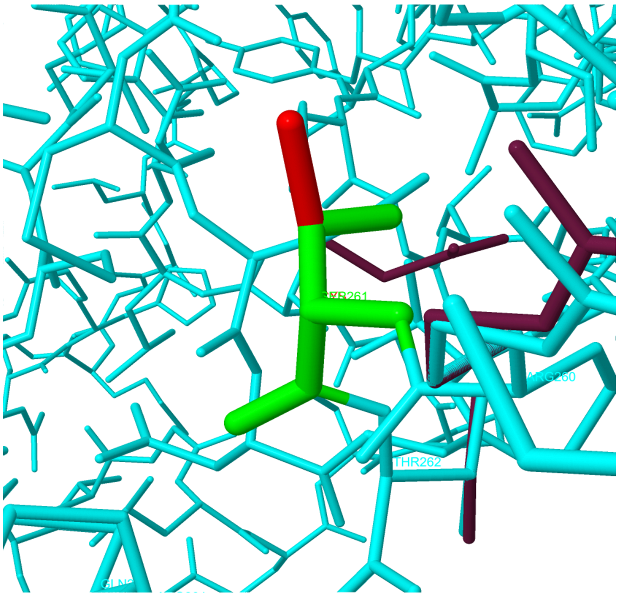

**Supplementary Figure 15:** Side chain replacement in S261C. This substitution leads to an expansion of cavity volume by 113.616 Å^3.

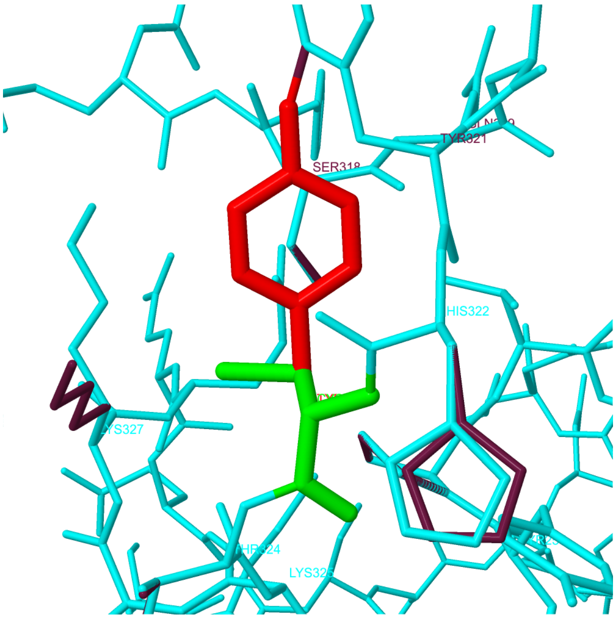

**Supplementary Figure 16:** Side chain replacement in C323Y. This substitution triggers a clash alert. The local clash score for the wild-type residue is 10.50 and the local clash score for the mutant residue is 40.58. The mutant structure has a MolProbity clash score ≥30 and the increase in clash score is >18 compared to the wild-type.

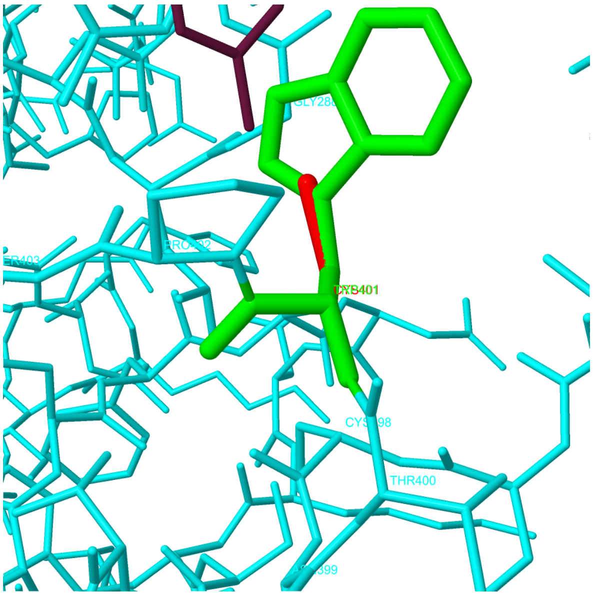

**Supplementary Figure 17:** Side chain replacement in W401C. This substitution leads to an expansion of cavity volume by 103.68 Å^3.

**Supplementary Table 1:** Deleterious SNPs predicted by PROVEAN, SIFT, and PolyPhen-2.

| dbSNP ID | Codon Change | AAS | PROVEAN | | SIFT | | PolyPhen-2 Prediction | |
|---|---|---|---|---|---|---|---|---|
| | | | Predic -tion | Score | Predic -tion | Score | Hum Div | Hum Var |
| rs1322050108 | C[T/C]C | L58P | Del | -5.86 | Dam | 0 | PD | PD |
| rs863223779 | A[C/A]G | T72K | Del | -5.31 | Dam | 0 | PD | PD |
| rs863223779 | A[C/T]G | T72M | Del | -5.31 | Dam | 0 | PD | PD |
| rs104894381 | [G/A]GA | G80R | Del | -7.29 | Dam | 0 | PD | PD |
| rs1131691460 | [T/C]TT | F84L | Del | -5.5 | Dam | 0 | PD | PD |
| rs1565941576 | C[C/T]C | P85L | Del | -9.16 | Dam | 0 | PD | PD |
| rs1565941579 | [C/T]CC | P85S | Del | -7.33 | Dam | 0 | PD | PD |
| rs775593443 | G[T/C]G | V91A | Del | -2.7 | Dam | 0.001 | PD | PD |
| rs770717773 | [G/A]GC | G93S | Del | -5.5 | Dam | 0.001 | PD | PD |
| rs890394778 | G[G/A]C | G93D | Del | -6.41 | Dam | 0.001 | PD | PD |
| rs890394778 | G[G/T]C | G93V | Del | -8.24 | Dam | 0 | PD | PD |
| rs1203503352 | [C/T]CC | P96S | Del | -3.75 | Dam | 0.024 | PD | PD |
| rs1463838783 | T[A/G]C | Y100C | Del | -8.24 | Dam | 0 | PD | PD |
| rs1463838783 | T[A/T]C | Y100F | Del | -3.66 | Dam | 0.004 | PD | PD |
| rs1350768465 | A[T/C]G | M104T | Del | -5.16 | Dam | 0.038 | PD | PD |
| rs1565941484 | [G/A]AC | D105N | Del | -4.59 | Dam | 0 | PD | PD |
| rs1369673380 | G[T/A]A | V107E | Del | -5.51 | Dam | 0 | PD | PD |
| rs376621016 | [C/A]CT | P108T | Del | -3.41 | Dam | 0.001 | PD | PD |
| rs1411518530 | GA[C/G] | D110E | Del | -3.47 | Dam | 0 | PD | PD |
| rs1411518530 | GA[C/A] | D110E | Del | -3.47 | Dam | 0 | PD | PD |
| rs1428277072 | [G/A]AC | D110N | Del | -4.19 | Dam | 0.001 | PD | PD |
| rs77357563 | [G/T]AT | D111Y | Del | -7.69 | Dam | 0 | PD | PD |
| rs483353129 | A[G/T]A | R113I | Del | -7.34 | Dam | 0 | PD | PD |
| rs483353129 | A[G/A]A | R113K | Del | -2.75 | Dam | 0 | PD | PD |
| rs757586102 | [T/C]AC | Y114H | Del | -4.59 | Dam | 0 | PD | PD |
| rs1269970792 | [T/C]GG | W121R | Del | -12.86 | Dam | 0 | PD | PD |
| rs863223773 | [G/C]GC | G125R | Del | -7.4 | Dam | 0 | PD | PD |
| rs1370517694 | [G/T]GC | G133C | Del | -4.87 | Dam | 0.023 | PD | PD |
| rs1057519050 | C[T/G]G | L135R | Del | -4.25 | Dam | 0 | PD | PD |
| rs1178440681 | G[T/C]G | V137A | Del | -3.7 | Dam | 0 | PD | PD |
| rs148715804 | GA[C/G] | D140E | Del | -3.7 | Dam | 0.008 | PD | PD |
| rs1424817435 | [C/T]CC | P142S | Del | -7.4 | Dam | 0 | PD | PD |
| rs374906778 | [G/A]CC | A143T | Del | -3.2 | Dam | 0.011 | PD | PD |
| rs1555226320 | [G/T]GG | G145W | Del | -7.4 | Dam | 0 | PD | PD |
| rs772248871 | G[T/A]C | V153D | Del | -6.29 | Dam | 0 | PD | PD |
| rs1464680326 | [C/T]TC | L160F | Del | -3.5 | Dam | 0.001 | PD | PD |
| rs1183256015 | C[A/T]T | H170L | Del | -4.92 | Dam | 0.001 | PD | PD |
| rs1565939347 | [A/G]AT | N174D | Del | -4.79 | Dam | 0.001 | PD | PD |
| rs1555226019 | G[T/G]G | V186G | Del | -6.42 | Dam | 0 | PD | PD |
| rs557183241 | [G/A]TG | V186M | Del | -2.71 | Dam | 0.006 | PD | PD |
| rs1392668757 | [G/T]AT | D189Y | Del | -2.81 | Dam | 0 | PD | PD |
| rs763004247 | [C/G]AC | H204D | Del | -5.74 | Dam | 0.002 | PD | PD |

| | | | | | | | | |
|---|---|---|---|---|---|---|---|---|
| rs190881877 | [G/T]TC | V205F | Del | -3.27 | Dam | 0.001 | PD | PoD |
| rs769113870 | GA[G/C] | E208D | Del | -2.78 | Dam | 0.001 | PD | PD |
| rs987371810 | [G/A]CA | A213T | Del | -3.33 | Dam | 0.039 | PD | PD |
| rs1318021626 | A[T/A]C | I222N | Del | -6.66 | Dam | 0 | PD | PD |
| rs1555225344 | A[C/T]G | T223M | Del | -5.66 | Dam | 0 | PD | PD |
| rs1565935426 | A[A/G]G | K226R | Del | -2.88 | Dam | 0 | PD | PD |
| rs1131691932 | [A/G]AG | K226E | Del | -3.84 | Dam | 0 | PD | PD |
| rs1565935405 | [T/G]TT | F232V | Del | -6.7 | Dam | 0 | PD | PD |
| rs104894378 | C[G/C]G | R237P | Del | -6.47 | Dam | 0 | PD | PD |
| rs104894378 | C[G/A]G | R237Q | Del | -3.76 | Dam | 0 | PD | PD |
| rs104894382 | [C/T]GG | R237W | Del | -7.29 | Dam | 0 | PD | PD |
| rs377625550 | [A/T]GC | S261C | Del | -2.61 | Dam | 0 | PD | PD |
| rs1298950685 | T[G/A]T | C323Y | Del | -2.95 | Dam | 0.001 | PD | PD |
| rs944423586 | C[C/A]C | P337H | Del | -2.99 | Dam | 0 | PD | PoD |
| rs377532269 | [C/T]GG | R375W | Del | -3.01 | Dam | 0 | PD | PD |
| rs377649723 | TG[G/T] | W401C | Del | -3.14 | Dam | 0 | PD | PD |

AAS= Amino Acid Substitution, Del=Deleterious, Dam=Damaging, PD=Probably Damaging & PoD=Possibly Damaging

**Supplementary Table 2:** Effects of amino acid substitutions on the TBX5 protein as predicted by MutPred2.

| AAS | Probability of deleterious effect | Molecular mechanisms with P-values <=0.05 | Probability | P-value |
|---|---|---|---|---|
| L58P | 0.951 | Altered Stability | 0.34 | 0.0042 |
| | | Loss of Helix | 0.29 | 0.01 |
| | | Gain of Strand | 0.28 | 0.01 |
| | | Altered Metal binding | 0.26 | 0.02 |
| T72K | 0.958 | Gain of Allosteric site at T72 | 0.34 | 0.00056 |
| | | Altered Metal binding | 0.33 | 0.0026 |
| | | Gain of Acetylation at T72 | 0.22 | 0.03 |
| | | Loss of Catalytic site at E73 | 0.17 | 0.02 |
| | | Altered DNA binding | 0.15 | 0.04 |
| T72M | 0.922 | Gain of Allosteric site at T72 | 0.29 | 0.0031 |
| | | Altered Metal binding | 0.27 | 0.0074 |
| | | Gain of Catalytic site at E73 | 0.16 | 0.02 |
| G80R | 0.968 | Gain of Allosteric site at G80 | 0.3 | 0.0016 |
| | | Altered DNA binding | 0.29 | 0.0036 |
| | | Loss of Strand | 0.27 | 0.02 |
| | | Loss of Relative solvent accessibility | 0.23 | 0.05 |
| | | Altered Disordered interface | 0.22 | 0.03 |
| | | Gain of Catalytic site at R81 | 0.18 | 0.01 |
| | | Loss of Methylation at K78 | 0.18 | 0.01 |
| | | Altered Metal binding | 0.14 | 0.02 |
| F84L | 0.923 | Altered Ordered interface | 0.35 | 0.0051 |
| | | Altered DNA binding | 0.28 | 0.0049 |
| | | Loss of Allosteric site at R82 | 0.26 | 0.01 |
| | | Loss of Acetylation at K88 | 0.23 | 0.02 |
| | | Loss of Catalytic site at R81 | 0.17 | 0.02 |
| | | Gain of Methylation at K88 | 0.1 | 0.04 |
| P85L | 0.887 | Altered Ordered interface | 0.32 | 0.01 |
| | | Loss of Allosteric site at R82 | 0.29 | 0.0051 |
| | | Altered DNA binding | 0.28 | 0.0048 |
| | | Loss of Acetylation at K88 | 0.25 | 0.01 |
| | | Loss of Relative solvent accessibility | 0.23 | 0.05 |
| | | Altered Disordered interface | 0.19 | 0.04 |
| | | Gain of Catalytic site at R81 | 0.19 | 0.01 |
| | | Gain of Methylation at K88 | 0.09 | 0.05 |
| P85S | 0.888 | Gain of Relative solvent accessibility | 0.27 | 0.02 |
| | | Loss of Allosteric site at R82 | 0.27 | 0.0084 |
| | | Altered DNA binding | 0.26 | 0.0058 |
| | | Gain of Acetylation at K88 | 0.24 | 0.02 |
| | | Altered Disordered interface | 0.22 | 0.03 |
| | | Loss of Catalytic site at R81 | 0.17 | 0.02 |
| V91A | 0.609 | Altered Ordered interface | 0.3 | 0.02 |
| | | Gain of Acetylation at K88 | 0.24 | 0.02 |

| | | | | |
|---|---|---|---|---|
| | | Altered DNA binding | 0.23 | 0.01 |
| | | Altered Stability | 0.16 | 0.02 |
| | | Gain of Methylation at K88 | 0.1 | 0.04 |
| G93S | 0.865 | Altered Ordered interface | 0.33 | 0.0074 |
| | | Gain of Relative solvent accessibility | 0.32 | 0.0038 |
| | | Altered DNA binding | 0.24 | 0.008 |
| | | Loss of Acetylation at K88 | 0.24 | 0.02 |
| G93D | 0.925 | Altered Ordered interface | 0.3 | 0.02 |
| | | Altered DNA binding | 0.3 | 0.0015 |
| | | Gain of Relative solvent accessibility | 0.27 | 0.02 |
| | | Loss of Strand | 0.26 | 0.03 |
| | | Loss of Acetylation at K88 | 0.25 | 0.01 |
| G93V | 0.927 | Loss of Relative solvent accessibility | 0.31 | 0.0072 |
| | | Altered Ordered interface | 0.29 | 0.02 |
| | | Loss of Acetylation at K88 | 0.25 | 0.01 |
| | | Altered DNA binding | 0.24 | 0.0093 |
| P96S | 0.806 | Altered Ordered interface | 0.33 | 0.0071 |
| | | Gain of Relative solvent accessibility | 0.33 | 0.0025 |
| | | Gain of Acetylation at K99 | 0.25 | 0.01 |
| | | Loss of Allosteric site at Y100 | 0.2 | 0.04 |
| | | Loss of Methylation at K99 | 0.1 | 0.04 |
| Y100C | 0.945 | Altered Ordered interface | 0.36 | 0.0044 |
| | | Loss of Acetylation at K99 | 0.31 | 0.0036 |
| | | Loss of Relative solvent accessibility | 0.28 | 0.02 |
| | | Altered Metal binding | 0.24 | 0.04 |
| | | Loss of Allosteric site at D105 | 0.24 | 0.02 |
| | | Gain of Catalytic site at D105 | 0.12 | 0.03 |
| | | Loss of Methylation at K99 | 0.11 | 0.04 |
| Y100F | 0.75 | Altered Ordered interface | 0.35 | 0.0052 |
| | | Loss of Relative solvent accessibility | 0.28 | 0.02 |
| | | Loss of Acetylation at K99 | 0.28 | 0.0063 |
| | | Altered Metal binding | 0.25 | 0.03 |
| | | Loss of Allosteric site at D105 | 0.24 | 0.02 |
| | | Gain of Catalytic site at D105 | 0.1 | 0.04 |
| | | Loss of Methylation at K99 | 0.1 | 0.04 |
| M104T | 0.945 | Altered Ordered interface | 0.3 | 0.02 |
| | | Gain of Relative solvent accessibility | 0.3 | 0.0086 |
| | | Altered Stability | 0.27 | 0.0067 |
| | | Altered Metal binding | 0.26 | 0.01 |
| | | Gain of Allosteric site at D105 | 0.24 | 0.01 |
| | | Gain of Acetylation at K99 | 0.24 | 0.02 |
| | | Gain of Methylation at K99 | 0.11 | 0.03 |
| | | Gain of Catalytic site at D105 | 0.1 | 0.04 |
| D105N | 0.913 | Altered Metal binding | 0.51 | 0.0036 |
| | | Altered Ordered interface | 0.29 | 0.03 |
| | | Gain of Strand | 0.26 | 0.04 |
| | | Gain of Allosteric site at D105 | 0.23 | 0.02 |
| | | Gain of Catalytic site at D105 | 0.1 | 0.04 |

| | | | | |
|---|---|---|---|---|
| V107E | 0.934 | Altered Metal binding | 0.42 | 0.00098 |
| | | Altered Ordered interface | 0.32 | 0.01 |
| | | Gain of Allosteric site at D105 | 0.24 | 0.01 |
| | | Gain of Catalytic site at D105 | 0.14 | 0.02 |
| P108T | 0.893 | Altered Metal binding | 0.29 | 0.005 |
| | | Altered Ordered interface | 0.28 | 0.04 |
| | | Loss of Allosteric site at D105 | 0.25 | 0.02 |
| | | Loss of Catalytic site at D105 | 0.1 | 0.04 |
| D110E | 0.822 | Altered Metal binding | 0.4 | 0.0012 |
| | | Altered Ordered interface | 0.24 | 0.04 |
| | | Gain of Allosteric site at D105 | 0.23 | 0.02 |
| | | Altered DNA binding | 0.17 | 0.03 |
| | | Gain of Catalytic site at D105 | 0.11 | 0.03 |
| D110N | 0.862 | Altered Metal binding | 0.47 | 0.0006 |
| | | Altered Ordered interface | 0.3 | 0.02 |
| | | Gain of Strand | 0.26 | 0.05 |
| | | Loss of Allosteric site at D105 | 0.22 | 0.03 |
| | | Altered DNA binding | 0.16 | 0.04 |
| | | Gain of Catalytic site at D105 | 0.11 | 0.04 |
| D111Y | 0.916 | Altered Metal binding | 0.54 | 0.00032 |
| | | Altered Ordered interface | 0.39 | 0.0011 |
| | | Gain of Strand | 0.27 | 0.02 |
| | | Gain of Allosteric site at I106 | 0.19 | 0.05 |
| | | Altered DNA binding | 0.17 | 0.04 |
| | | Gain of Sulfation at D111 | 0.03 | 0.01 |
| R113I | 0.937 | Gain of Relative solvent accessibility | 0.36 | 0.0011 |
| | | Altered Metal binding | 0.33 | 0.001 |
| | | Altered Ordered interface | 0.3 | 0.02 |
| | | Gain of Acetylation at K115 | 0.19 | 0.05 |
| | | Altered DNA binding | 0.17 | 0.04 |
| | | Gain of Methylation at K115 | 0.09 | 0.05 |
| R113K | 0.894 | Altered Metal binding | 0.33 | 0.0038 |
| | | Gain of Relative solvent accessibility | 0.29 | 0.01 |
| | | Gain of Acetylation at R113 | 0.26 | 0.0083 |
| | | Altered Ordered interface | 0.24 | 0.04 |
| | | Altered DNA binding | 0.17 | 0.04 |
| | | Gain of Methylation at K115 | 0.09 | 0.04 |
| Y114H | 0.933 | Altered Metal binding | 0.41 | 0.0021 |
| | | Altered Ordered interface | 0.31 | 0.01 |
| | | Loss of Relative solvent accessibility | 0.27 | 0.02 |
| | | Altered DNA binding | 0.17 | 0.03 |
| | | Altered Stability | 0.13 | 0.03 |
| W121R | 0.965 | Gain of Intrinsic disorder | 0.3 | 0.04 |
| | | Loss of Relative solvent accessibility | 0.28 | 0.02 |
| | | Loss of Strand | 0.26 | 0.05 |
| | | Loss of Allosteric site at W121 | 0.21 | 0.04 |
| | | Gain of Methylation at K126 | 0.16 | 0.01 |
| G125R | 0.944 | Gain of Allosteric site at W121 | 0.24 | 0.02 |

| | | | | |
|---|---|---|---|---|
| | | Loss of Methylation at K126 | 0.12 | 0.03 |
| G133C | 0.763 | Gain of Allosteric site at R134 | 0.34 | 0.00046 |
| | | Altered Metal binding | 0.27 | 0.0072 |
| | | Gain of Catalytic site at H138 | 0.2 | 0.01 |
| L135R | 0.952 | Altered Metal binding | 0.44 | 0.00082 |
| | | Altered Stability | 0.43 | 0.0026 |
| | | Loss of Loop | 0.3 | 0.0059 |
| | | Gain of Allosteric site at R134 | 0.25 | 0.01 |
| | | Loss of Catalytic site at D140 | 0.22 | 0.0084 |
| V137A | 0.697 | Altered Stability | 0.41 | 0.0028 |
| | | Loss of Allosteric site at R134 | 0.26 | 0.01 |
| | | Altered Metal binding | 0.24 | 0.01 |
| | | Loss of Catalytic site at D140 | 0.21 | 0.01 |
| D140E | 0.744 | Altered Metal binding | 0.3 | 0.0054 |
| | | Gain of Catalytic site at D140 | 0.24 | 0.0053 |
| | | Loss of Allosteric site at H138 | 0.23 | 0.03 |
| P142S | 0.747 | Loss of Loop | 0.27 | 0.03 |
| | | Altered Metal binding | 0.26 | 0.0093 |
| | | Loss of Allosteric site at H138 | 0.24 | 0.02 |
| | | Gain of Catalytic site at D140 | 0.21 | 0.0094 |
| A143T | 0.541 | Altered Metal binding | 0.24 | 0.01 |
| | | Loss of Allosteric site at H138 | 0.23 | 0.02 |
| | | Loss of Catalytic site at D140 | 0.21 | 0.01 |
| G145W | 0.921 | Loss of Loop | 0.27 | 0.04 |
| | | Altered Metal binding | 0.25 | 0.01 |
| | | Altered Ordered interface | 0.25 | 0.02 |
| | | Gain of Catalytic site at D140 | 0.23 | 0.0065 |
| V153D | 0.942 | | | |
| L160F | 0.833 | Gain of Strand | 0.26 | 0.04 |
| | | Gain of Allosteric site at N163 | 0.19 | 0.05 |
| | | Loss of Catalytic site at K159 | 0.09 | 0.05 |
| H170L | 0.936 | Altered Ordered interface | 0.29 | 0.03 |
| | | Altered Metal binding | 0.28 | 0.02 |
| | | Loss of Allosteric site at H170 | 0.19 | 0.05 |
| N174D | 0.741 | Altered DNA binding | 0.29 | 0.0023 |
| | | Altered Ordered interface | 0.27 | 0.05 |
| | | Altered Metal binding | 0.25 | 0.0093 |
| | | Loss of Allosteric site at H177 | 0.25 | 0.02 |
| V186G | 0.898 | Altered Stability | 1 | 0.00012 |
| | | Gain of Intrinsic disorder | 0.31 | 0.04 |
| | | Altered Metal binding | 0.22 | 0.02 |
| V186M | 0.709 | Altered Metal binding | 0.22 | 0.02 |
| | | Altered Stability | 0.18 | 0.01 |
| D189Y | 0.823 | Altered Metal binding | 0.23 | 0.02 |
| H204D | 0.902 | Altered Ordered interface | 0.29 | 0.02 |
| | | Gain of Loop | 0.27 | 0.03 |
| | | Altered Transmembrane protein | 0.15 | 0.02 |
| V205F | 0.855 | Altered Transmembrane protein | 0.13 | 0.02 |

| | | | | |
|---|---|---|---|---|
| E208D | 0.809 | Altered Transmembrane protein | 0.14 | 0.02 |
| A213T | 0.805 | Altered Ordered interface | 0.25 | 0.02 |
| | | Altered Transmembrane protein | 0.21 | 0.0043 |
| | | Altered DNA binding | 0.16 | 0.04 |
| | | Loss of Catalytic site at Y217 | 0.15 | 0.02 |
| | | Altered Metal binding | 0.13 | 0.03 |
| T223M | 0.724 | Altered Ordered interface | 0.33 | 0.008 |
| | | Gain of Helix | 0.27 | 0.04 |
| | | Altered Metal binding | 0.22 | 0.02 |
| | | Altered Transmembrane protein | 0.22 | 0.0032 |
| | | Altered DNA binding | 0.17 | 0.03 |
| I222N | 0.943 | Altered Ordered interface | 0.32 | 0.0091 |
| | | Altered Metal binding | 0.24 | 0.02 |
| | | Altered Transmembrane protein | 0.21 | 0.0045 |
| | | Gain of Catalytic site at Y217 | 0.18 | 0.01 |
| | | Altered DNA binding | 0.18 | 0.03 |
| K226R | 0.435 | | | |
| K226E | 0.726 | Loss of Helix | 0.27 | 0.04 |
| | | Altered Transmembrane protein | 0.23 | 0.0029 |
| F232V | 0.754 | Altered Disordered interface | 0.37 | 0.0063 |
| | | Loss of Acetylation at K234 | 0.29 | 0.0051 |
| | | Altered DNA binding | 0.17 | 0.03 |
| | | Altered Transmembrane protein | 0.12 | 0.03 |
| | | Gain of Proteolytic cleavage at R237 | 0.12 | 0.03 |
| R237P | 0.937 | Altered Disordered interface | 0.29 | 0.03 |
| | | Gain of Loop | 0.29 | 0.0068 |
| | | Loss of Acetylation at K234 | 0.25 | 0.01 |
| | | Altered DNA binding | 0.15 | 0.04 |
| | | Loss of Proteolytic cleavage at R237 | 0.13 | 0.02 |
| R237Q | 0.716 | Loss of Acetylation at K234 | 0.25 | 0.01 |
| | | Altered DNA binding | 0.15 | 0.04 |
| | | Loss of Proteolytic cleavage at R237 | 0.13 | 0.03 |
| R237W | 0.834 | Loss of Intrinsic disorder | 0.38 | 0.03 |
| | | Gain of Loop | 0.27 | 0.02 |
| | | Loss of Acetylation at K234 | 0.25 | 0.01 |
| | | Altered DNA binding | 0.2 | 0.02 |
| | | Loss of Proteolytic cleavage at R237 | 0.13 | 0.03 |
| S261C | 0.5 | Loss of ADP-ribosylation at R260 | 0.23 | 0.02 |
| | | Loss of SUMOylation at K266 | 0.2 | 0.04 |
| | | Loss of O-linked glycosylation at T262 | 0.14 | 0.04 |
| | | Loss of Proteolytic cleavage at R260 | 0.11 | 0.04 |
| C323Y | 0.623 | Gain of Phosphorylation at Y321 | 0.3 | 0.01 |
| | | Loss of SUMOylation at K327 | 0.26 | 0.0069 |
| | | Gain of Sulfation at Y321 | 0.03 | 0.01 |
| P337H | 0.255 | | | |
| R375W | 0.676 | Loss of Phosphorylation at Y380 | 0.31 | 0.02 |
| | | Loss of Pyrrolidone carboxylic acid at Q376 | 0.06 | 0.03 |
| W401C | 0.76 | Gain of Intrinsic disorder | 0.45 | 0.0036 |

| | | | | |
|---|---|---|---|---|
| | | Gain of Loop | 0.27 | 0.03 |
| | | Loss of Disulfide linkage at C398 | 0.1 | 0.05 |

**Supplementary Table 3:** Functional SNPs on the 3′ UTR of *TBX5* gene predicted by MirSNP.

| Sl. No. | Gene | miRNA | SNP | Effect | Allele | Score | Energy | Conservation |
|---|---|---|---|---|---|---|---|---|
| 1 | *TBX5* | hsa-miR-1256 | rs184793497 | break | C | 140 | -9.41 | 0.006 |
| | | | | | A | | | |
| 2 | *TBX5* | hsa-miR-1294 | rs76633287 | break | T | 156 | -20.21 | 0.052 |
| | | | | | C | | | |
| 3 | *TBX5* | hsa-miR-130a-5p | rs181130921 | decrease | G | 146 | -11.65 | 0.499 |
| | | | | | C | 145 | -7.2 | 0.499 |
| 4 | *TBX5* | hsa-miR-130b-5p | rs10850326 | decrease | G | 158 | -16.41 | 0.004 |
| | | | | | A | 148 | -11.31 | 0.007 |
| 5 | *TBX5* | hsa-miR-130b-5p | rs112808902 | decrease | - | 158 | -16.41 | 0 |
| | | | | | AAGAGA | 148 | -11.31 | 0.002 |
| 6 | *TBX5* | hsa-miR-1323 | rs140532076 | enhance | T | 151 | -13 | 0 |
| | | | | | C | 155 | -14.92 | 0 |
| 7 | *TBX5* | hsa-miR-141-3p | rs112051831 | break | G | 154 | -14.34 | 0 |
| | | | | | A | | | |
| 8 | *TBX5* | hsa-miR-144-5p | rs188337192 | break | T | 155 | -15.19 | 0 |
| | | | | | C | | | |
| 9 | *TBX5* | hsa-miR-144-5p | rs192445542 | break | T | 155 | -15.19 | 0 |
| | | | | | C | | | |
| 10 | *TBX5* | hsa-miR-147a | rs184793497 | create | C | | | |
| | | | | | A | 149 | -14.29 | 0.004 |
| 11 | *TBX5* | hsa-miR-200a-3p | rs112051831 | break | G | 153 | -12.15 | 0 |
| | | | | | A | | | |
| 12 | *TBX5* | hsa-miR-204-5p | rs10850326 | break | G | 147 | -16.09 | 0.002 |
| | | | | | A | | | |
| 13 | *TBX5* | hsa-miR-21-3p | rs12426660 | enhance | T | 150 | -17.2 | 0 |
| | | | | | C | 154 | -18.97 | 0 |
| 14 | *TBX5* | hsa-miR-211-3p | rs12426660 | break | T | 150 | -19.42 | 0.002 |
| | | | | | C | | | |
| 15 | *TBX5* | hsa-miR-211-5p | rs10850326 | break | G | 147 | -14.74 | 0.002 |
| | | | | | A | | | |
| 16 | *TBX5* | hsa-miR-2355-3p | rs76694710 | create | C | | | |
| | | | | | A | 160 | -17.09 | 0.052 |
| 17 | *TBX5* | hsa-miR-23a-3p | rs181130921 | decrease | G | 147 | -9.06 | 0.499 |
| | | | | | C | 144 | -8.22 | 0.499 |

| | | | | | | | | |
|---|---|---|---|---|---|---|---|---|
| 18 | *TBX5* | hsa-miR-2467-3p | rs28730760 | create | A | | | |
| | | | | | C | 143 | -16 | 0.005 |
| 19 | *TBX5* | hsa-miR-29b-1-5p | rs181130921 | break | G | 159 | -23.67 | 0.237 |
| | | | | | C | | | |
| 20 | *TBX5* | hsa-miR-3119 | rs184360838 | create | G | | | |
| | | | | | A | 140 | -10.91 | 0.023 |
| 21 | *TBX5* | hsa-miR-3134 | rs2384410 | break | T | 140 | -11.18 | 0.02 |
| | | | | | C | | | |
| 22 | *TBX5* | hsa-miR-3148 | rs75334514 | decrease | T | 146 | -8.63 | 0.004 |
| | | | | | A | 145 | -7.44 | 0.004 |
| 23 | *TBX5* | hsa-miR-3148 | rs192527148 | decrease | T | 148 | -7.26 | 0.423 |
| | | | | | G | 146 | -8.63 | 0.423 |
| 24 | *TBX5* | hsa-miR-3154 | rs143511878 | break | T | 163 | -27.56 | 0.853 |
| | | | | | C | | | |
| 25 | *TBX5* | hsa-miR-3160-5p | rs184360838 | break | G | 169 | -28.28 | 0.033 |
| | | | | | A | | | |
| 26 | *TBX5* | hsa-miR-3170 | rs76347803 | create | G | | | |
| | | | | | C | 150 | -16.34 | 0.019 |
| 27 | *TBX5* | hsa-miR-3170 | rs76633287 | create | T | | | |
| | | | | | C | 150 | -16.34 | 0.015 |
| 28 | *TBX5* | hsa-miR-3170 | rs78828439 | create | C | | | |
| | | | | | A | 150 | -16.34 | 0.02 |
| 29 | *TBX5* | hsa-miR-3180-5p | rs143511878 | break | T | 146 | -19.83 | 0.298 |
| | | | | | C | | | |
| 30 | *TBX5* | hsa-miR-3183 | rs10850326 | enhance | G | 145 | -21.31 | 0.018 |
| | | | | | A | 147 | -21.26 | 0.018 |
| 31 | *TBX5* | hsa-miR-3183 | rs112808902 | enhance | - | 146 | -19 | 0.042 |
| | | | | | AAGAGA | 147 | -21.26 | 0 |
| 32 | *TBX5* | hsa-miR-3189-3p | rs76694710 | create | C | | | |
| | | | | | A | 151 | -19.19 | 0.018 |
| 33 | *TBX5* | hsa-miR-3202 | rs143511878 | create | T | | | |
| | | | | | C | 151 | -19.87 | 0.853 |
| 34 | *TBX5* | hsa-miR-330-3p | rs143511878 | decrease | T | 145 | -15.27 | 0.878 |
| | | | | | C | 144 | -14.11 | 0.878 |
| 35 | *TBX5* | hsa-miR-340-3p | rs28730761 | create | A | | | |
| | | | | | G | 148 | -16.52 | 0.254 |
| 36 | *TBX5* | hsa-miR-3613-3p | rs147774482 | decrease | T | 164 | -10.03 | 0.999 |
| | | | | | C | 156 | -10.05 | 0.999 |
| 37 | *TBX5* | hsa-miR-3678-3p | rs28730760 | create | A | | | |
| | | | | | C | 146 | -15.86 | 0.005 |
| 38 | *TBX5* | hsa-miR-374c-3p | rs183595902 | break | T | 149 | -17.08 | 0.139 |
| | | | | | G | | | |
| 39 | *TBX5* | hsa-miR-4257 | rs28730760 | create | A | | | |
| | | | | | C | 156 | -18.19 | 0.005 |
| 40 | *TBX5* | hsa-miR-4282 | rs75334514 | create | T | | | |
| | | | | | A | 140 | -2.08 | 0 |
| 41 | *TBX5* | hsa-miR-4286 | rs183595902 | break | T | 144 | -15.71 | 0.466 |
| | | | | | G | | | |

| | | | | | | | | |
|---|---|---|---|---|---|---|---|---|
| 42 | *TBX5* | hsa-miR-4287 | rs10850326 | break | G | 140 | -18.5 | 0.001 |
| | | | | | A | | | |
| 43 | *TBX5* | hsa-miR-4311 | rs187379531 | decrease | G | 148 | -9.94 | 0.001 |
| | | | | | C | 146 | -10.85 | 0.001 |
| 44 | *TBX5* | hsa-miR-4427 | rs117414057 | break | T | 150 | -10.75 | 0.004 |
| | | | | | C | | | |
| 45 | *TBX5* | hsa-miR-4446-3p | rs117414057 | enhance | T | 157 | -20.94 | 0.006 |
| | | | | | C | 161 | -22.89 | 0.006 |
| 46 | *TBX5* | hsa-miR-4455 | rs2384410 | create | T | | | |
| | | | | | C | 148 | -16.7 | 0.002 |
| 47 | *TBX5* | hsa-miR-4635 | rs76799455 | decrease | T | 165 | -25.41 | 0.005 |
| | | | | | A | 157 | -23.67 | 0.005 |
| 48 | *TBX5* | hsa-miR-4647 | rs117414057 | enhance | T | 161 | -20.78 | 0.095 |
| | | | | | C | 165 | -23.09 | 0.095 |
| 49 | *TBX5* | hsa-miR-4656 | rs117414057 | enhance | T | 159 | -26.31 | 0.006 |
| | | | | | C | 163 | -27.88 | 0.006 |
| 50 | *TBX5* | hsa-miR-4685-3p | rs10850326 | break | G | 147 | -17.89 | 0.001 |
| | | | | | A | | | |
| 51 | *TBX5* | hsa-miR-4688 | rs12426660 | create | T | | | |
| | | | | | C | 145 | -21.04 | 0.005 |
| 52 | *TBX5* | hsa-miR-4697-3p | rs148346089 | create | T | | | |
| | | | | | C | 160 | -28.56 | 0.024 |
| 53 | *TBX5* | hsa-miR-4711-5p | rs148346089 | break | T | 143 | -19.27 | 0 |
| | | | | | C | | | |
| 54 | *TBX5* | hsa-miR-4717-3p | rs116382074 | break | G | 148 | -15.92 | 0.03 |
| | | | | | A | | | |
| 55 | *TBX5* | hsa-miR-4723-3p | rs10850326 | decrease | G | 155 | -22.13 | 0.018 |
| | | | | | A | 150 | -20.84 | 0.018 |
| 56 | *TBX5* | hsa-miR-4724-5p | rs140532076 | break | T | 145 | -13.87 | 0 |
| | | | | | C | | | |
| 57 | *TBX5* | hsa-miR-4753-3p | rs10850326 | enhance | G | 154 | -16.7 | 0.004 |
| | | | | | A | 166 | -22.9 | 0.002 |
| 58 | *TBX5* | hsa-miR-4753-3p | rs112808902 | enhance | - | 154 | -16.7 | 0 |
| | | | | | AAGAGA | 166 | -22.9 | 0.001 |
| 59 | *TBX5* | hsa-miR-4764-5p | rs2384410 | break | T | 156 | -20.66 | 0.002 |
| | | | | | C | | | |
| 60 | *TBX5* | hsa-miR-4771 | rs12426660 | create | T | | | |
| | | | | | C | 160 | -17.33 | 0.23 |
| 61 | *TBX5* | hsa-miR-486-3p | rs12426660 | create | T | | | |
| | | | | | C | 150 | -20.67 | 0.01 |
| 62 | *TBX5* | hsa-miR-494 | rs116382074 | decrease | G | 157 | -15.31 | 0.697 |
| | | | | | A | 149 | -10.83 | 0.697 |
| 63 | *TBX5* | hsa-miR-499a-3p | rs116382074 | break | G | 153 | -21.11 | 0.271 |
| | | | | | A | | | |
| 64 | *TBX5* | hsa-miR-499b-3p | rs116382074 | break | G | 146 | -12.79 | 0.271 |
| | | | | | A | | | |
| 65 | *TBX5* | hsa-miR-5006-5p | rs148346089 | decrease | T | 151 | -22.42 | 0.001 |
| | | | | | C | 146 | -23.1 | 0.001 |

| | | | | | | | | |
|---|---|---|---|---|---|---|---|---|
| 66 | *TBX5* | hsa-miR-5007-3p | rs116382074 | create | G | | | |
| | | | | | A | 140 | -8.37 | 0.066 |
| 67 | *TBX5* | hsa-miR-5010-3p | rs184793497 | decrease | C | 150 | -9.37 | 0 |
| | | | | | A | 140 | -5.98 | 0.007 |
| 68 | *TBX5* | hsa-miR-5093 | rs78058633 | create | G | | | |
| | | | | | A | 149 | -14.21 | 0 |
| 69 | *TBX5* | hsa-miR-5094 | rs148346089 | break | T | 152 | -16.17 | 0.024 |
| | | | | | C | | | |
| 70 | *TBX5* | hsa-miR-513a-3p | rs75334514 | create | T | | | |
| | | | | | A | 153 | -8.8 | 0 |
| 71 | *TBX5* | hsa-miR-513c-3p | rs75334514 | create | T | | | |
| | | | | | A | 153 | -5.75 | 0 |
| 72 | *TBX5* | hsa-miR-5197-5p | rs187379531 | create | G | | | |
| | | | | | C | 152 | -15.54 | 0.002 |
| 73 | *TBX5* | hsa-miR-545-5p | rs141223729 | enhance | T | 157 | -12.28 | 1 |
| | | | | | C | 161 | -14.27 | 1 |
| 74 | *TBX5* | hsa-miR-548a-3p | rs140532076 | enhance | T | 147 | -13.58 | 0 |
| | | | | | C | 151 | -15.87 | 0 |
| 75 | *TBX5* | hsa-miR-548an | rs147774482 | create | T | | | |
| | | | | | C | 146 | -12.7 | 0.909 |
| 76 | *TBX5* | hsa-miR-548e | rs140532076 | enhance | T | 154 | -9.62 | 0 |
| | | | | | C | 158 | -11.97 | 0 |
| 77 | *TBX5* | hsa-miR-548f | rs140532076 | decrease | T | 145 | -7.01 | 0 |
| | | | | | C | 142 | -6.35 | 0 |
| 78 | *TBX5* | hsa-miR-548o-3p | rs140532076 | enhance | T | 149 | -9.53 | 0 |
| | | | | | C | 150 | -11.15 | 0 |
| 79 | *TBX5* | hsa-miR-5572 | rs76694710 | create | C | | | |
| | | | | | A | 143 | -19.81 | 0.028 |
| 80 | *TBX5* | hsa-miR-5584-5p | rs76799455 | break | T | 154 | -13.2 | 0.003 |
| | | | | | A | | | |
| 81 | *TBX5* | hsa-miR-576-3p | rs28730760 | break | A | 159 | -18.42 | 0.003 |
| | | | | | C | | | |
| 82 | *TBX5* | hsa-miR-576-3p | rs112051831 | enhance | G | 155 | -18.63 | 0 |
| | | | | | A | 159 | -18.42 | 0 |
| 83 | *TBX5* | hsa-miR-599 | rs6489956 | break | G | 145 | -9.33 | 0.001 |
| | | | | | A | | | |
| 84 | *TBX5* | hsa-miR-616-3p | rs188337192 | create | T | | | |
| | | | | | C | 157 | -15.51 | 0.001 |
| 85 | *TBX5* | hsa-miR-624-5p | rs141223729 | enhance | T | 157 | -19.6 | 1 |
| | | | | | C | 161 | -21.59 | 1 |
| 86 | *TBX5* | hsa-miR-944 | rs192527148 | break | T | 158 | -8.44 | 0 |
| | | | | | G | | | |

**Supplementary Table 4:** 3D protein model evaluations by PROCHECK, Verify 3D, and SWISS-MODEL Structure Assessment Tool.

| Model | PROCHECK | | | | Verify 3D (% residues with an average 3D-1D score >=0.2) | MolProbity Score | QMEAN Z-score |
|---|---|---|---|---|---|---|---|
| | Number of residues in most favored regions | Number of residues in additional allowed regions | Number of residues in generously allowed regions | Number of residues in disallowed regions | | | |
| I-TASSER | 364 (84.1%) | 55 (12.7%) | 3 (0.7%) | 11 (2.5%) | 69.69 | 1.03 | -4.68 |
| Phyre2 | 342 (79.0%) | 78 (18.0%) | 6 (1.4%) | 7 (1.6%) | 70.66 | 1.17 | -7.05 |
| Robetta | 409 (94.5%) | 23 (5.3%) | 0 (0.0%) | 1 (0.2%) | 70.46 | 0.5 | -1.37 |
